# The conserved function of bHLH121 and overexpression of N-terminal bHLH121 fragment suppresses iron deficiency response in *Arabidopsis*

**DOI:** 10.64898/2026.09.27.754851

**Authors:** Peijun Zhou, Chuanfa Liu, Yilin Pan, Yuchen Fei, Renfang Shen, Ruonan Wang, Ping Lan

**Affiliations:** State Key Laboratory of Soil and Sustainable Agriculture, Institute of Soil Science, Chinese Academy of Sciences, Nanjing 211135, China; University of Chinese Academy of Sciences, Beijing 100049, China; University of Chinese Academy of Sciences, Nanjing, 211135, China

**Keywords:** bHLH121, function conservation, fragment, overexpression, suppress, Fe deficiency response, *Arabidopsis*

## Abstract

Iron (Fe) homeostasis is fundamental to plant growth and development and is centrally regulated by a well-conserved network of basic helix-loop-helix (bHLH) transcription factors. In *Arabidopsis thaliana*, bHLH121 regulates the expression of *FER-like iron deficiency-induced transcription factor* (*FIT*) and clade Ib *bHLH38/39/100/101* genes, among others. However, its functional conservation remains unexplored, and some contradictory results have been reported regarding the effects of *bHLH121* overexpression. Here, we investigated the evolutionary conservation of bHLH121 through complementary tests in the *Arabidopsis bhlh121* mutant created by CRISPR/Cas9 gene editing. We also investigated the effects of ectopically expressing full-length, N-terminal, and C-terminal fragments of *bHLH121* on plant responses to Fe deficiency in *Arabidopsis*. Transgenic plants overexpressing the full-length *bHLH121* driven by the cauliflower mosaic virus 35S promoter (*bHLH121 ox*) resulted in shorter roots and more severe leaf chlorosis compared to wild-type plants under Fe-limiting conditions. Moreover, overexpression of the N-terminal fragment (*N121 ox*) rendered plants highly sensitive to Fe deficiency, whereas overexpression of the C-terminal fragment (*C121 ox*) did not. Consistent with the phenotypic observations, *N121 ox* plants accumulated lower Fe content, and the ferric chelate reductase activity was significantly reduced. In *N121 ox* plants, the induction of key Fe regulatory genes, especially *FIT*, was significantly inhibited under Fe deficiency. Confocal imaging showed that both N121-GFP and C121-GFP fusion proteins could localize to the nucleus. Electrophoretic mobility shift assay showed that both full-length bHLH121 and N121 could effectively bind the probes derived from the promoter of *FIT*. The present work reveals a conserved function of bHLH121 in Fe homeostasis across several dicot species and demonstrates that the N121 may possess a dominant nature in Fe deficiency responses.

## 1. Introduction

Iron (Fe) is an essential micronutrient for nearly all organisms, serving as a redox-active cofactor in critical physiological processes such as photosynthesis, respiration, and hormone biosynthesis (Liang, 2022; Marschner, 1995). Despite its geological abundance, Fe primarily exists as insoluble Fe(III) oxides in neutral to alkaline soils, limiting its bioavailability and making it a common growth-limiting factor that impacts crop yield and quality. Conversely, excess Fe can trigger ROS production via the Fenton reaction, posing toxicity risks. To maintain homeostasis, plants tightly regulate Fe uptake and allocation. Two distinct acquisition strategies have evolved (Romheld & Marschner, 1986): non-graminaceous plants like *Arabidopsis thaliana* use Strategy I, where proton extrusion (*Arabidopsis* plasma membrane H⁺-ATPase isoform 2, AHA2) (Santi & Schmidt, 2009), coumarin secretion (Pleiotropic drug resistance 9, PDR9) (Robe et al., 2021), Fe(III) reduction (Ferric reductase oxidase 2, FRO2) (Robinson et al., 1999), and Fe(II) transport (Iron-regulated transporter 1, IRT1) (Eide et al., 1996; Vert et al., 2002) facilitate Fe solubilization and uptake, while graminaceous species employ Strategy II, secreting mugineic acid-family phytosiderophores to directly chelate Fe(III) (Bashir et al., 2006; Curie et al., 2001; Nozoye et al., 2011). However, this strict dichotomy has recently been challenged, as non-graminaceous plants have also been shown to utilize Fe(III)-chelating mechanisms similar to Strategy II (Robe et al., 2025).

Fe deficiency induces the expression of genes involved in Fe acquisition and translocation, which are rigorously controlled at the transcriptional level by specialized transcription factors (TFs) (Liang, 2022). Among these regulators, basic helix-loop-helix (bHLH) family members have emerged as central players in maintaining Fe homeostasis. The bHLH TF FER-like iron deficiency-induced transcription factor (FIT) serves as a master regulator of Fe uptake in *Arabidopsis*, where it forms functional complexes with clade Ib bHLH proteins (bHLH38, bHLH39, bHLH100, and bHLH101) to activate downstream targets including *IRT1* and *FRO2* (Colangelo & Guerinot, 2004; Yuan et al., 2008). Notably, these interactions not only enable transcriptional activation but also stabilize FIT protein, which otherwise undergoes ubiquitin-mediated degradation via the RING E3 ligases BRUTUS-like 1(BTSL1) and BTSL2 and 26S proteasome (Rodríguez-Celma et al., 2019). Both *FIT* and clade Ib *bHLH* genes are Fe-deficiency inducible and positively regulated by clade IVc bHLH TFs (bHLH34, bHLH104, bHLH105/IAA-leucine resistant 3 (ILR3), and bHLH115) (Li et al., 2016; Liang et al., 2017; Zhang et al., 2015). Intriguingly, while clade IVc bHLH genes maintain constitutive expression under all Fe conditions, their protein products— particularly bHLH105/ILR3 and bHLH115—are post-translationally regulated through BTS- and BTSL1/2-mediated ubiquitination and proteasomal degradation (Hindt et al., 2017; Selote et al., 2015; Zhao et al., 2026), mirroring the regulatory mechanism observed for FIT.

Recent studies have identified bHLH121/Upstream Regulator of IRT1 (URI), a clade IVb bHLH transcription factor, as a pivotal regulator of Fe deficiency response (Gao et al., 2020; Kim et al., 2019; Lei et al., 2020). Unlike Fe-responsive genes, *bHLH121* overall exhibits constitutive expression throughout the plant regardless of Fe status. However, Fe availability influences bHLH121 cellular localization in roots (Gao et al., 2020), most likely through protein phosphorylation modification (Kim et al., 2019) and its interaction with clade IVc bHLH TFs (Lei et al., 2020). Genetic evidence demonstrates that *bhlh121* loss-of-function mutants develop severe Fe deficiency symptoms, accompanied by impaired induction of key Fe-regulated genes (*bHLH38/39/100/101*). This regulatory role of bHLH121 in the Fe deficiency response appears to be conserved, as its homologs in other species have been implicated in similar functions: in petunia, *PhbHLH121* has been characterized as an important regulator of Fe deficiency tolerance (Pan et al., 2024), while in rice, *OsbHLH064* functions as a central regulator of Fe homeostasis and enhances grain Fe accumulation (Gao et al., 2026). Furthermore, a recent phloem sap proteomic study revealed that bHLH121 may regulate the production or loading of potential long-distance signaling factors that coordinate root Fe uptake with shoot Fe status (Nathalie et al., 2025). Mechanistically, the regulation of *FIT* by bHLH121 remains controversial. While one study reported that bHLH121 directly binds the *FIT* promoter and synergistically enhances *FIT* activation by clade IVc bHLH TFs (Lei et al., 2020), two other major studies did not observe in planta binding of bHLH121 to the *FIT* promoter (Gao et al., 2020; Kim et al., 2019). Although the functional conservation of the transcriptional regulatory network involving FIT and clade Ib bHLH genes has been demonstrated (Grillet & Schmidt, 2019), direct evidence to support that bHLH121 is functionally conserved across different plant species is lacking. Notably, conflicting overexpression phenotypes reported for different bHLH121 isoforms (Gao et al., 2020; Lei et al., 2020) suggest context-dependent regulation.

In this study, we first created loss-of-function *bhlh121* mutants by CRISPR-Cas9 method. Then we investigated the evolutionary conservation of bHLH121 function in Fe homeostasis across several dicotyledonous species. Furthermore, we employed ectopic overexpression of distinct bHLH121 protein variants in *Arabidopsis* to elucidate its segment-specific contributions to Fe homeostasis regulation. In agreement with previous findings (Lei et al., 2020), constitutive 35S-driven expression of full-length *bHLH121* conferred heightened sensitivity to Fe deficiency, as evidenced by significantly shorter primary roots and more severe leaf chlorosis relative to wild-type plants. Intriguingly, transgenic lines expressing the N-terminal fragment (*N121 ox*) exhibited suppressed Fe deficiency response, whereas those expressing the C-terminal fragment (*C121 ox*) did not. Transcriptional characterization revealed that overexpression of different bHLH121 forms specifically regulates several key genes in Fe homeostasis, including *FIT* and *bHLH38/39*, under Fe-deficient conditions. Furthermore, biochemical analyses suggest that N121 competes with full-length bHLH121 for target gene binding.

## 2. Materials and methods

### 2.1. Plant materials and growth conditions

*Arabidopsis thaliana* ecotype Columbia (Col-0) plants were used as the wild-type (WT) and served as the genetic background for transgenic plants. The *bhlh121-7* and *bhlh121-8* mutants were generated in this study using the CRISPR/Cas9 method with the *pEn-C1.1* (pEN) and *pDe-CAS9-D10A* (pDE) vectors (Schiml et al., 2014) as previously described (Xue et al., 2021). Oligos used for CRISPR/Cas9-mediated bHLH121 mutation are listed in Supplementary Table S2. *Arabidopsis* seeds were sterilized by first washing them with 75% alcohol for 3 minutes, then with a 0.5% (v/v) NaClO solution containing 0.5% (v/v) Tween-20 for 10 minutes, and finally rinsing them with sterile deionized water five times. After being stratified at 4 °C for 2 days, the sterilized seeds were sown on ES medium and placed in a greenhouse with 22 ℃/16-h-light and 20 ℃/8-h-dark photoperiod for culture. The ES medium (Estelle & Somerville, 1987) was used as the control treatment that contained 5 mM KNO_3_, 2 mM Ca(NO_3_)_2_⋅4H_2_O, 2 mM MgSO_4_⋅7H_2_O, 2.5 mM KH_2_PO_4_, 40 μM Fe(III)-EDTA, 14 μM MnCl_2_⋅4H_2_O, 70 μM H_3_BO_3_, 1 μM ZnSO_4_⋅7H_2_O, 0.5 μM CuSO_4_⋅5H_2_O, 0.2 μM NaMoO_4_⋅2H_2_O, and 10 μM NaCl. The complete ES medium was used for the +Fe treatment. The ES medium without Fe(III)-EDTA was used for the 0Fe treatment. The Fe chelator ferrozine [3-(2-pyridyl)-5,6-diphenyl-1,2,4-triazine sulfonate] was added at 100 µM to the Fe-free ES medium for the -Fe treatment as described previously (Lan et al., 2012).

The growth media for various nutrient deficiency treatments were prepared by modifying the standard ES medium. For the low K treatment, KNO_3_ was omitted and replaced with KCl at a final concentration of 10 µM, while NH_4_NO_3_ was added to maintain nitrogen levels. For the -P treatment, KH_2_PO_4_ was excluded and KCl was supplemented to keep potassium concentrations constant. For the -Zn and -Cu treatments, ZnSO_4_ or CuSO_4_ were omitted respectively, with K_2_SO_4_ added to balance sulfate levels. For the -Mn treatment, MnCl_2_ was removed and replaced with KCl to maintain chloride concentration. All media were supplemented with 1% (w/v) sucrose, 0.1% (w/v) MES, and 0.8% (w/v) agar, and adjusted to pH 5.5.

### 2.2 Cloning of bHLH121 homologous genes

Total RNA was extracted from the leaves of poplar (*Populus trichocarpa*), cucumber (Jinyan No. 4), soybean (Zhonghuang 37), and citrus (Shatangju) using TRIzol reagent (Vazyme), and then reverse transcribed into cDNA using the PrimeScript RT reagent Kit with gDNA Eraser (TaKaRa, RP047A). Specific primers containing restriction enzyme sites were designed based on the cDNA sequences of *bHLH121* homologs *PtbHLH121* (Potri.004G168100), *CsbHLH121* (KGN61896), *GsbHLH121* (XP_028243911), and *CcbHLH121* (ESR36836). Primers are listed in Supplementary Table S2.

### 2.3 Construction of vectors and generation of transgenic plants

To obtain *probHLH121::bHLH121:GFP* plasmid for complementation lines, the full-length genome sequence of *bHLH121* (excluding the stop codon and 3’ UTR; TAIR10 transcript ID: At3g19860.2) was amplified and ligated into the modified binary vector *pCAMBIA1300-GFP* in which the CaMV 35S promoter was replaced by the native promoter of *bHLH121*, a region of 3000 bp upstream from the initiation codon of *bHLH121*. To obtain *bHLH121* overexpression lines, the full-length coding region of *bHLH121*, without 5’/3’ UTR, was amplified from Col-0 cDNA using primers based on the TAIR10 transcript model At3g19860.2, which encodes a 337-amino-acid protein, and cloned into the modified binary vector *pCAMBIA1300-GFP* (Xue et al., 2021), which was driven by the cauliflower mosaic virus *CaMV-35S* promoter (*pro35S::bHLH121:GFP*). Using the sequence-verified plasmid (*pro35S::bHLH121:GFP*) as template, *N121* (amino acids 1–160 of bHLH121) and *C121* (amino acids 151–337 of bHLH121) fragments were cloned and subsequently ligated into the modified binary vector *pCAMBIA1300-GFP* (*pro35S::N121:GFP*, *pro35S::C121:GFP*). For ectopic expression of *bHLH121* homologs from four dicotyledonous species, the cDNA sequences of *bHLH121* homologous genes were cloned by PCR amplification with the high-fidelity enzyme PrimeSTAR (TaKaRa), and individually ligated the correctly sequenced cDNA sequences into the modified binary vector *pCAMBIA1300-GFP* to obtain recombinant plasmids.

All these recombinant plasmids were transformed into *Agrobacterium* strain GV3101. Subsequently, the wild-type *Arabidopsis* (WT) (for *pro35S::bHLH121:GFP*, *pro35S::N121:GFP*, and *pro35S::C121:GFP*) and the *bhlh121-7* mutant (for *probHLH121::bHLH121:GFP*) were transformed using the floral dipping method (Clough & Bent, 1998). T0 transgenic seeds were screened on 1/2 MS medium containing 25 mg/L hygromycin to obtain transgenic lines. T3 homozygous seedlings were used for further analyses. Primers used in these experiments are listed in Table S2.

### 2.4 Measurement of root length and fresh weight

*Arabidopsis* seeds were sown directly onto the medium and cultivated vertically in a growth chamber for 10 days. Phenotypic observations and photographic documentation were subsequently conducted. Root length was measured using ImageJ software (National Institutes of Health, USA). Fresh weight was determined by weighing seedlings in batches of ten, and the results are showed as the average fresh weight per seedling.

### 2.5 Measurement of chlorophyll content

The shoots of *Arabidopsis* were collected and extracted in 80% acetone in the dark until the samples were decolorized completely. The supernatant was taken and the absorbance (A) at 645 and 663 nm was then measured using a spectrophotometer (SpectraMax Plus 384 Absorbance Microplate Reader, Molecular Devices). The total chlorophyll content was calculated using the following formula: chlorophyll content = (20.2 × A645 + 8.02 × A663) × V/FW and was expressed as micrograms per gram fresh weight (Arnon, 1949).

### 2.6 Bioinformatic analysis of protein amino acid sequences

Using the amino acid sequence of *Arabidopsis bHLH121* as the input sequence, the homologous proteins of *bHLH121* in poplar (*Populus trichocarpa*), cucumber (*Cucumis sativus*), citrus (*Citrus clementina*), soybean (*Glycine soja*), carrot (*Daucus carota*), red bean (*Vigna angularis*), rose (*Rosa chinensis*), coffee (*Coffea canephora*), cassava (*Manihot esculenta*) were screened out through BLAST in the Ensembl Plants database (http://plants.ensembl.org/index.html). These species were chosen to represent diverse dicot lineages, including woody perennials (poplar, citrus), legumes (soybean, red bean), and herbaceous crops (cucumber, carrot, cassava), to assess the breadth of bHLH121 functional conservation. The phylogenetic tree was constructed by MEGA 7.0.21 (https://www.Megasoftware.net/) using the neighbor-joining method with 1000 bootstrap replicates. MEME (https://meme-suite.org/meme/tools/meme) was used to identify conserved motifs of amino acid sequence. The specific setting parameters were as follows: 6 < motif width < 50, 10 motifs were identified. Multiple sequence alignment analysis was performed using the DNAMAN software (v8).

The basic physicochemical properties of the protein, including the number of amino acids, theoretical isoelectric point, molecular weight, instability index, aliphatic index, and grand average of hydropathicity (GRAVY), were predicted using the ProtParam (Wilkins et al., 1999). Nuclear localization signals were identified with NLStradamus (Nguyen Ba et al., 2009). The bHLH domain was identified with InterPro (Blum et al., 2025). The L-ZIP domain is recognized by the canonical heptad repeat motif L-X_6_-L-X_6_-L-X_6_-L, in which “L” refers to leucine and “X” stands for any amino acid residue. Intrinsically disordered regions (IDRs) of bHLH121 were predicted via the PONDR (http://www.pondr.com/) (Linding et al., 2003). The tertiary structure of the protein was modeled using the SWISS-MODEL (Waterhouse et al., 2018).

### 2.7 Fe(III) chelate reductase assay

Fe(III) chelate reductase activity was determined as previously described with minor modifications (Yi & Guerinot, 1996). Plants were grown for 7 days on ES medium and transferred to +Fe or -Fe growth conditions for 5 days. For qualitative analysis, five seedlings per line were grouped and placed on chromogenic medium containing 250 μM Fe(III)-EDTA, 250 μM Ferrozine, 400 μM CaSO_4_, and 0.7% agar. The plates were incubated in darkness for 12 h and then photographed as described previously (Bauer, 2016). For quantitative analysis, plant roots were immersed in a chromogenic solution consisting of 300 µM Ferrozine and 100 µM Fe(III)-EDTA, followed by incubation at room temperature in darkness for 30 min. Absorbance at 562 nm was measured, and root ferric-chelate reductase activity was calculated according to the following formula, reductase activity (μmol Fe(Ⅱ) gram^-1^h^-1^) = [A562/0.0286/0.5] × 0.001 / root weight (gram) / 0.5 (hour).

### 2.8 Determination of iron content

Iron content was measured using the bathophenanthroline disulfonate (BPDS) colorimetric method as previously described (Gautam et al., 2021) with minor modifications. Briefly, whole seedlings (roots and shoots combined) were used for Fe content measurement, samples were thoroughly washed three times with ultrapure water to remove surface-associated Fe, and three independent biological replicates were included for each measurement, with each replicate consisting of approximately 50 mg of well-dried plant tissue. Each sample was digested with 2 mL of 65% (v/v) nitric acid and 1 mL of 30% (v/v) hydrogen peroxide at 150 °C. The digested samples were mixed with an assay buffer containing 1 mM BPDS, 0.6 M sodium acetate, and 0.48 M hydroxylamine hydrochloride. Following incubation at room temperature in the dark for 30 min, the absorbance of the reaction mixture was measured at 535 nm using a microplate spectrophotometer. Iron concentration was determined by reference to a standard curve prepared with FeCl_3_ standards in the same assay solution and normalized to sample dry weight.

### 2.9 Subcellular localization observation

Fluorescence images were taken from roots using a confocal laser scanning microscope (STELLARIS 5, Leica). The overexpression lines were grown on +Fe or 0Fe conditions for 5 days before imaging. Excitation was performed at 488 nm, with emission signals collected between 505 and 530 nm for GFP fluorescence. GFP fluorescence intensity and nuclear to cytoplasmic fluorescence ratio were analyzed quantitatively using LAS X software. More than six transgenic lines (at least five roots for each line) were observed for each construct.

### 2.10 Quantitative real-time PCR analysis

Total RNA was extracted using TRIzol reagent (Vazyme) according to the manufacturer’ s instructions. 1 μg of RNA was used and reverse transcribed into cDNA using a PrimeScript™ RT reagent Kit with gDNA Eraser (TaKaRa). The cDNA, which was diluted 12-fold with RNase-free water, served as a template for the real-time fluorescent quantitative PCR reaction. The RT-qPCR assays were performed using a SYBR® Premix Ex Taq™ II (Tli RNaseH Plus) (TaKaRa) kit under a PIKOREAL 96 Real-Time PCR System (ThermoFisher Scientific). The thermal cycle regimes were as follows: 95 °C for 7 min, followed by 40 cycles of 95 °C for 5 s, and 60 °C for 30 s. *Tubulin alpha-3* (*TUA3*, At5g19770) was used as a reference gene. The relative expression was calculated using the formula 2^-△△CT^ (Livak & Schmittgen, 2001). Primers used are listed in Table S2.

### 2.11 Electrophoretic mobility shift assay (EMSA)

EMSAs were conducted using the Chemiluminescent EMSA Kit (Beyotime, China) following the manufacturer’s protocol. The full-length *bHLH121*, *N121*, and *C121* fragments were cloned into the pCold™ TF DNA vector, which adds a ∼52 kDa N-terminal Trigger Factor (TF) solubility tag with 6xHis-tag. The recombinant His-TF-bHLH121, His-TF-N121 and His-TF-C121 proteins were expressed by cold induction in *E. coli* and purified using BeyoGold™ His-tag Purification Resin under non-denaturing conditions. The DNA fragments of the *FIT* were synthesized and biotin was added to the 5’ terminus of the DNA. Unlabeled fragments of the same sequences or mutated sequences were used as competitors, and the His-tagged TF (His-TF) protein purified from *E. coli* transformed with the empty vector was used as the negative control. Probes used in these assays are listed in Table S2.

### 2.12 Statistical analysis

All data are represented as mean ± standard deviation (SD). GraphPad Prism 9.5 software was utilized to perform the statistical analysis and produce the graphs. For multiple comparisons of means, one-way ANOVA was performed followed by Tukey’s HSD test (*P* < 0.05). Student’s t-test (*P* < 0.05) was conducted for comparisons of means between two groups. There are at least 3 independent biological replicates for each experiment.

## 3. Results

### 3.1 CRISPR-Cas9-mediated generation of bHLH121 loss-of-function mutants in Arabidopsis

To investigate the functional conservation of *bHLH121*, we first generated CRISPR-Cas9-mediated knockout mutants in the wild-type Col-0 (WT) background, designated as *bhlh121-7* and *bhlh121-8* to distinguish them from previously reported *bhlh121* mutants (Gao et al., 2020; Lei et al., 2020). Both mutants are null alleles with premature termination of bHLH121 protein translation and show lower expression levels than WT (Fig. S1A-C). They also exhibit severe defects in the induction of Fe deficiency-responsive genes, including *IRT1*, *FRO2*, *bHLH38* and *bHLH39* under Fe depletion (Fig. S1D), suggesting that both mutants are loss-of-function *bhlh121* alleles. In agreement with previous studies, both *bhlh121* mutants showed hypersensitivity to Fe deficiency, as evidenced by severe leaf chlorosis and significantly reduced root elongation (Fig. 1A). Since both mutants exhibited comparable phenotypes, we reasoned that they were genetically similar, and selected *bhlh121-7* as a representative mutant for subsequent analyses. As expected, a native promoter-driven *bHLH121-GFP* fusion construct *ProbHLH121::bHLH121:GFP* successfully restored root growth, biomass, and chlorophyll content in the *bhlh121-7* mutant under Fe deficiency (Fig. 1B-D), further validating the functionality of bHLH121-GFP and confirming that *bHLH121* loss is causal for the Fe deficiency-sensitive phenotypes. Similarly, abolished Fe chelate reductase activity under Fe deficiency in the *bhlh121-7* mutant was also substantially rescued by bHLH121-GFP (Fig. 1E, F). Notably, those complemented lines with the expression levels of *bHLHI21-GFP* comparable to the endogenous *bHLH121* level in WT showed best restoration and a negative effect was caused by higher *bHLH121-GFP* expression level, especially in root growth (Fig. 1), indicating that the expression pattern of endogenous bHLH121 is tightly controlled for Fe homeostasis.

**Figure 1.**
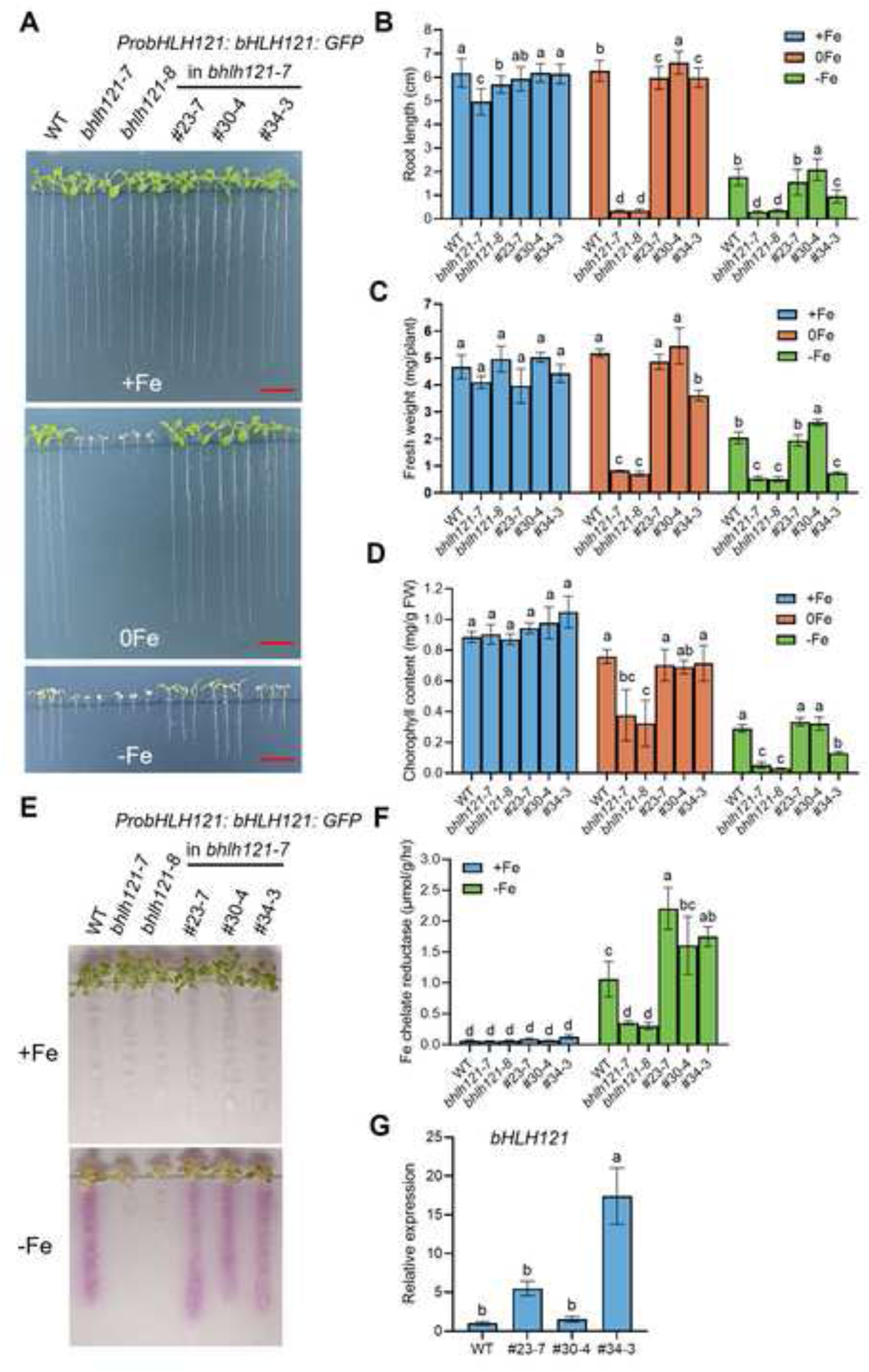
The *bhlh121* mutants exhibit high sensitivity to iron deficiency. (**A**) Phenotypes of the wild-type (WT), two *bhlh121* mutants (*bhlh121-7*, *bhlh121-8*) and three independent *bhlh121-7* lines complemented with the *ProbHLH121::bHLH121:GFP* transgene (#23-7, #30-4, #34-3) grown on ES medium with +Fe (40 μM Fe), 0Fe (0 μM Fe) or –Fe (0 μM Fe + 100 μM Ferrozine) for 10 days. Scale = 1 cm. (**B**) Root length (n = 30), (**C**) fresh weight (3n = 30), and (**D**) chlorophyll content (n = 3) of the WT, two *bhlh121* mutants and three complemented lines grown on +Fe, 0Fe, and –Fe medium for 10 d. (**E, F**) Qualitative (**E**) and quantitative (**F**) detection of ferric chelate reductase activity in the WT, two *bhlh121* mutants and three complemented lines. Seedlings grown on ES medium for 7 days were transferred to +Fe/-Fe medium for 5 days, and root tissues were used for the detection of ferric chelate reductase activity (n = 3). (**G**) Relative expression of *bHLH121* in three complemented lines. 7-day-old seedlings cultured on ES medium were harvested for qRT-PCR (n = 3). Data are means ± SD. Different letters represent significant differences determined by one-way ANOVA with Tukey’s post hoc test (*P* < 0.05).

### 3.2 Conserved function of bHLH121 homologs for Fe homeostasis regulation in several dicotyledonous species

An important question raised in this study is whether the function of bHLH121 is conserved across different plant species, which is crucial for evaluating its potential as a target for crop genetic improvement. To this end, we performed amino acid sequence alignment analysis. Using the *Arabidopsis* bHLH121 amino acid sequence as the query, we obtained the amino acid sequences of bHLH121 homologs from nine dicotyledonous species (*Populus trichocarpa*, *Manihot esculenta*, *Rosa chinensis*, *Citrus clementina*, *Vigna angularis*, *Glycine max*, *Daucus carota*, *Coffea canephora*, and *Cucumis sativus*) through BLAST searches against the Ensembl Plants database. Phylogenetic and conserved motif analysis of these sequences identified two bHLH transcription factor subfamilies corresponding to IVb and IVc (Fig. S2). Based on this, we speculated that the functions of these putative bHLH121 homologs from IVb subfamily may be conserved. Of course, it should be noted that a more comprehensive phylogenetic analysis incorporating all clade IVb and IVc members across species is required to fully resolve the evolutionary landscape of bHLH121 homologs.

To functionally verify the potential evolutionary conservation, we cloned four bHLH121 homologous genes from poplar, soybean, cucumber, and citrus, and performed heterologous expression in the *bhlh121-7* mutant for functional complementation assays. Remarkably, the homologs from all four species rescued both the root and shoot growth defect phenotypes of the *Arabidopsis bhlh121-7* mutant to varying degrees when grown on +Fe, 0Fe, or -Fe medium for 10 days (Figs. 2A, B, S3). And the complementation efficacy was significantly correlated with the expression level of each homologous gene in the tested transgenic lines (Fig. 2C). Due to the severe retarded plant growth on -Fe media, we measured the Fe contents of the plants grown under both control (+Fe) and 0Fe conditions. The results showed that Fe accumulation in transgenic plants expressing different *bHLH121* homologous genes was restored to levels comparable to those of WT plants (Fig. 3), indicating Fe deficiency response is rescued by expression of these homologous genes in *bhlh121* mutant.

**Figure 2.**
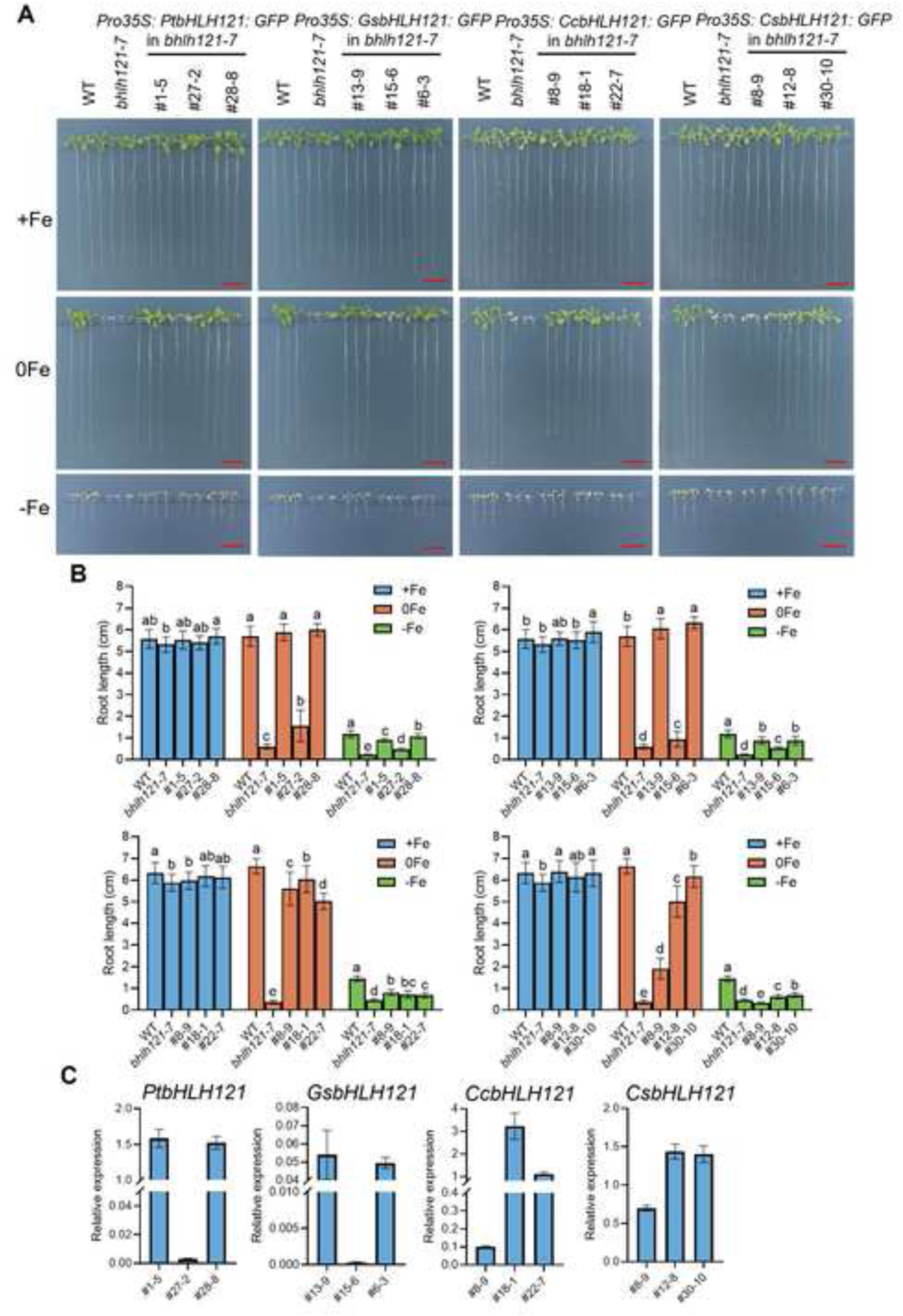
Overexpression of *bHLH121* homologous genes partially rescues the iron deficiency tolerance of the *bhlh121* mutant. (**A**) Phenotypes of the *bHLH121* homologous gene (*PtbHLH121*, *GsbHLH121*, *CcbHLH121*, *CsbHLH121*) complemented lines grown on +Fe, 0Fe, and -Fe medium for 10 days. Scale bar = 1 cm. (**B**) Root length (n = 24) of the *bHLH121* homologous gene complemented lines grown on +Fe, 0Fe, and -Fe medium for 10 d. (**C**) Relative expression of *PtbHLH121, GsbHLH121, CcbHLH121, CsbHLH121* in the *bHLH121* homologous gene complemented lines. Data are means ± SD. Different letters represent significant differences determined by one-way ANOVA with Tukey’s post hoc test (*P* < 0.05).

**Figure 3.**
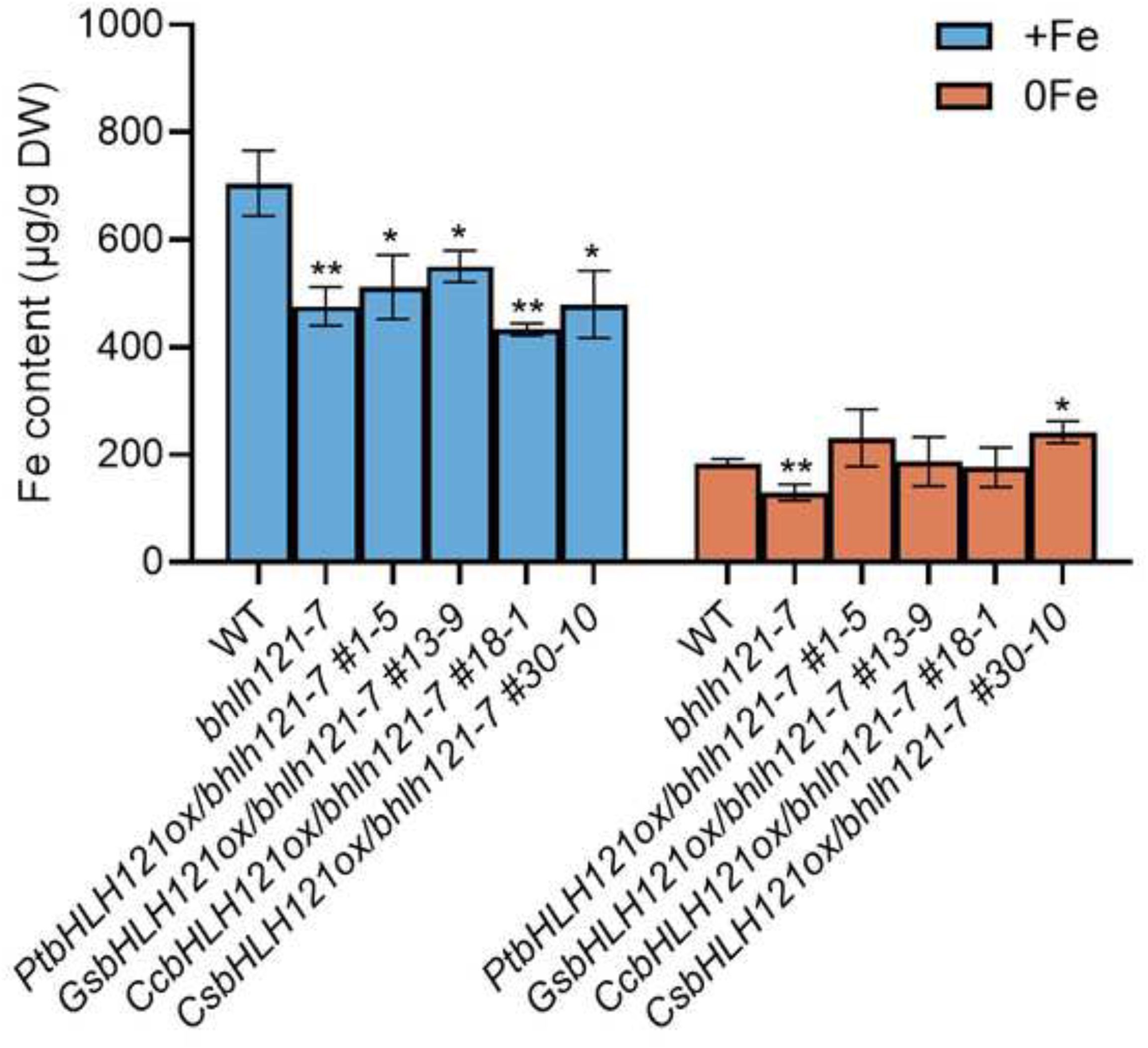
Overexpression of *bHLH121* homologous genes enhances the iron accumulation of the *bhlh121* mutant under Fe deficiency. Fe contents of the WT, the *bhlh121* mutant and the *bHLH121* homologous gene complementary lines grown on +Fe or 0Fe medium for 10 d. Data are means ± SD (n = 3). The unpaired two-tailed Student’s t-test was used to assess the statistical significance of the differences from the WT (\**P* < 0.05, and \*\**P* < 0.01).

To further validate the molecular mechanism underlying this putative functional conservation, we also performed assays of root ferric chelate reductase activity (FCR) and examined the expression of Fe deficiency-responsive genes. For these assays, seedlings were first grown on ES medium for 7 days, then transferred to +Fe or -Fe medium for an additional 5 days. Compared to WT control, expression of *PtbHLH121*, *GsbHLH121*, and *CcbHLH121* significantly restored FCR activity (Fig. 4A, B). Unexpectedly, little or no obvious FCR was detected in *bhlh121* mutant expressing *CsbHLH121*, presumably due to the drastically reduced induction of *IRT1* and *FRO2* under -Fe conditions (Fig. 4), which suggests a possible functional divergence among bHLH121 homologs across plant species. Altogether, these results suggest that bHLH121’s function in regulating Fe homeostasis is overall conserved in most dicots, highlighting the potential of leveraging bHLH121 homologs to modulate Fe utilization across a broad spectrum of dicotyledonous crops.

**Figure 4.**
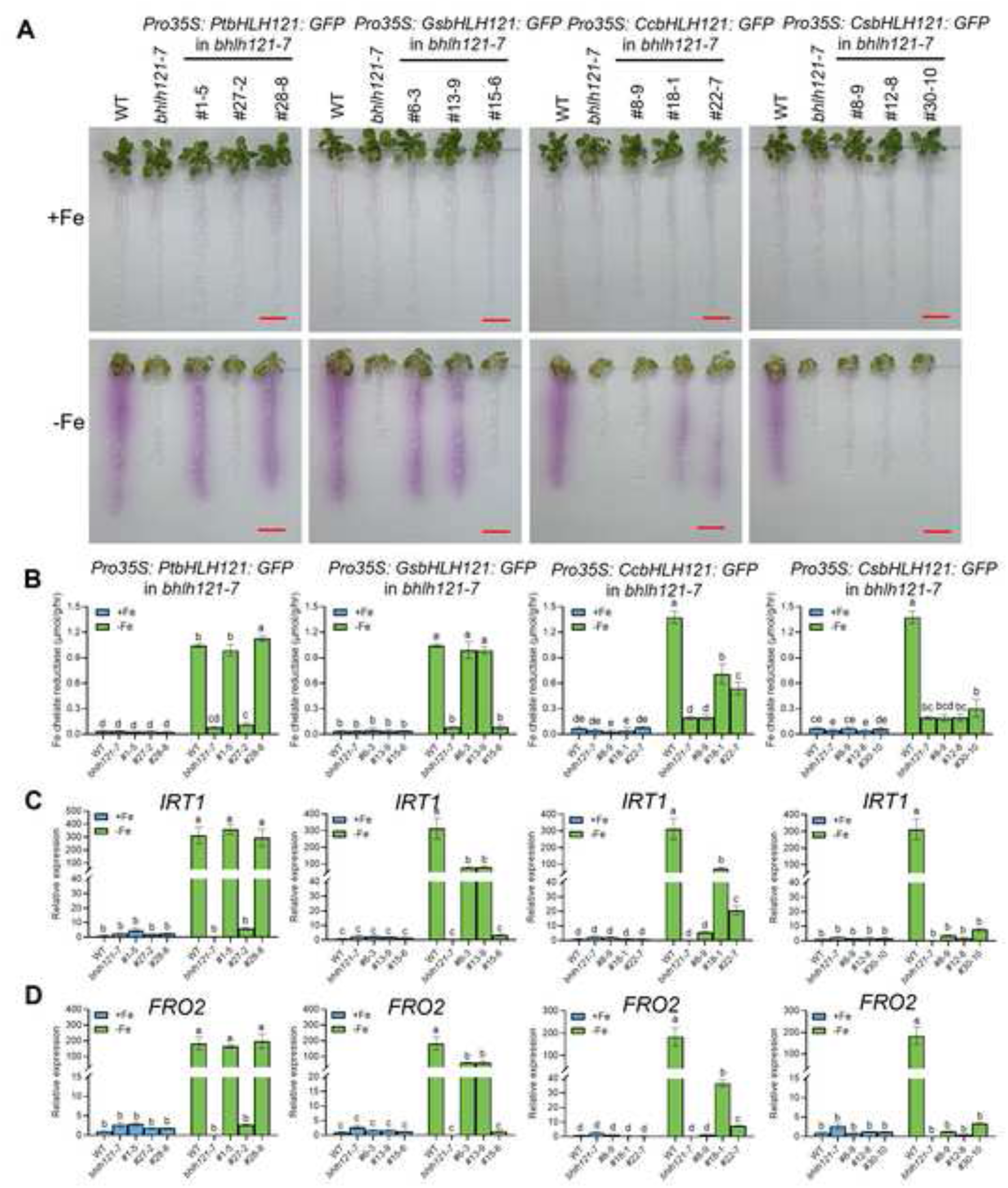
Overexpression of *bHLH121* homologous genes partially restores the iron deficiency response of the *bhlh121* mutant. (**A**, **B**) Qualitative (**A**) and quantitative (**B**) detection of ferric chelate reductase activity in the *bHLH121* homologous gene complemented lines. (**C**, **D**) Relative expression of *IRT1* (**C**) *and FRO2* (**D**) in the *bHLH121* homologous gene complemented lines. Seedlings grown on ES medium for 7 days were transferred to +Fe/-Fe medium for 5 days, and root tissues were used for the detection of ferric chelate reductase activity and gene expression. Data are means ± SD (n = 3). Different letters represent significant differences determined by one-way ANOVA with Tukey’s post hoc test (*P* < 0.05).

### 3.3 Overexpression of bHLH121 significantly inhibited root elongation under Fe deficiency

Our results demonstrated that higher expression of *bHLH121-GFP* showed negative effect on the restoration of the defective phenotypes in the *bhlh121-7* mutant under Fe deficiency (Fig. 1). The overexpression effect of *bHLH121* gene exhibited significant discrepancies in several aspects between the two research groups (Gao et al., 2020; Lei et al., 2020). To clarify this, we performed independent *bHLH121* overexpression assays in WT background. More than 10 independent stable transgenic lines (*bHLH121 ox*) were obtained for testing the response to Fe deficiency. While most lines were shown phenotypes similar to WT plants under Fe-replete conditions, the majority displayed more severe growth retardation under Fe deficiency, characterized by shortened primary roots and exacerbated leaf chlorosis (Fig. S4A-C). Notably, Pearson correlation analysis revealed a significant negative correlation between *bHLH121* transcript abundance and root growth under Fe deprivation conditions (Fig. S4D, E). The root length of the *bHLH121 ox* lines was negatively correlated with *bHLH121* expression levels when grown in ES medium without Fe but supplemented with 100 μM ferrozine, with a correlation coefficient (r) of −0.7117.

To further confirm this phenomenon, we measured root length, biomass, and chlorophyll content in three representative *bHLH121 ox* lines under Fe deficiency. The results showed that all three physiological parameters were negatively correlated with *bHLH121* expression levels (Fig. 5). Specifically, line #6-2, which harbors WT-comparable bHLH121 transcript level (Fig. 5B), displayed slightly reduced root growth inhibition under Fe deficiency (Fig. 5A, C). Conversely, line #15-5 with the highest *bHLH121* level exhibited the most defective growth phenotypes (Fig. 5). Taken together, consistent with our previous descriptions (Fig. 1), these results clearly demonstrate that overexpression of *bHLH121* significantly inhibited plant growth and root elongation under Fe deficiency.

**Figure 5.**
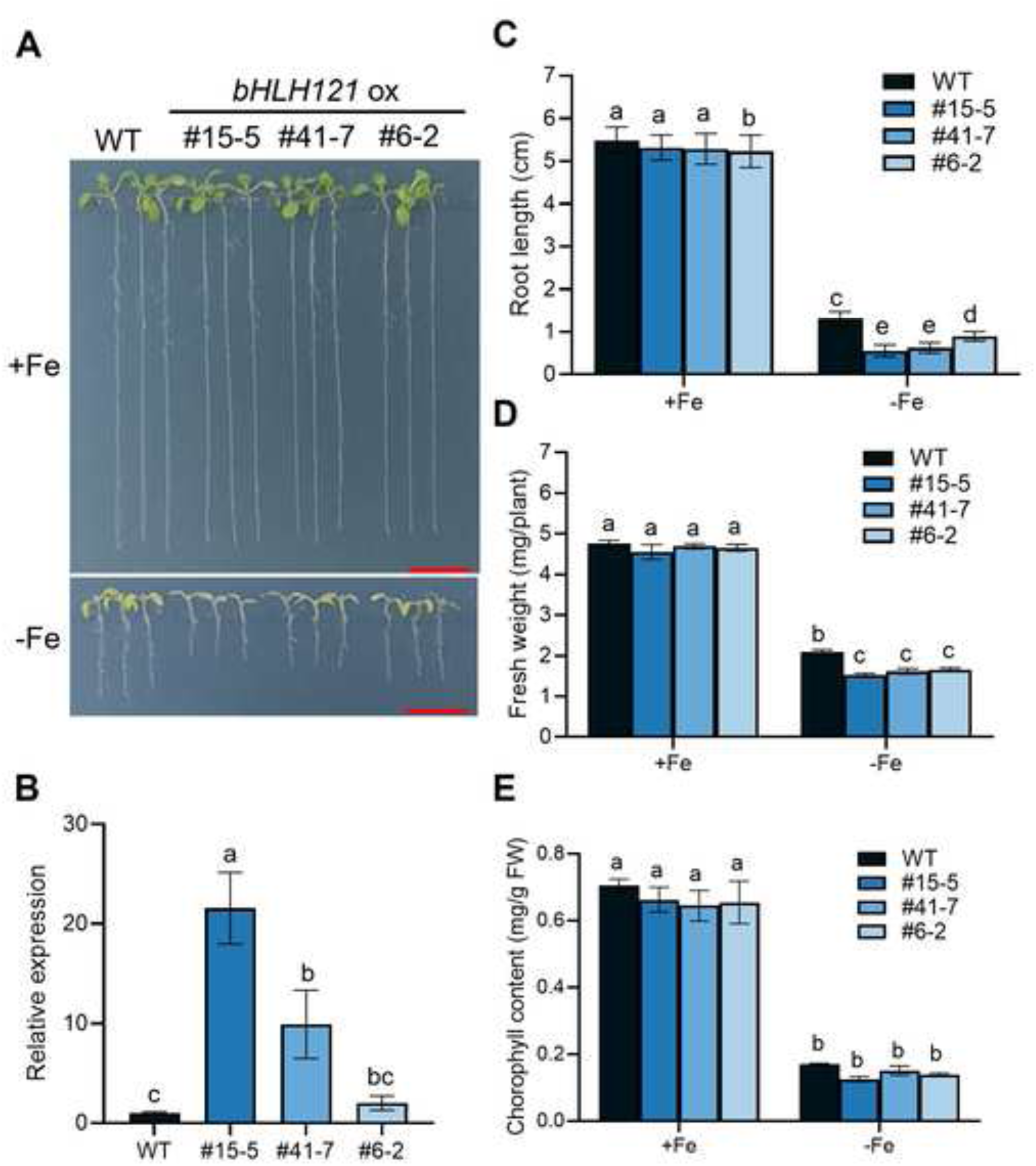
Overexpression of *bHLH121* significantly inhibited root elongation under iron deficiency. (**A**) Phenotypes of 10-day-old seedlings grown on +Fe and –Fe medium. Scale bars = 1 cm. (**B**) Gene expression levels of *bHLH121* in representative transgenic lines. (**C**) Root length analysis (n = 30). (**D**) Seedling fresh weight analysis (3n = 30). (**E**) Chlorophyll contents analysis (n = 3). Data are means ± SD. Different letters represent significant differences determined by one-way ANOVA with Tukey’s post hoc test (*P* < 0.05).

### 3.4 Sequence and structure analysis of bHLH121 protein in Arabidopsis

Following the observation that full-length overexpression of the *bHLH121* leads to Fe-deficiency hypersensitive phenotype, we sought to investigate the underlying molecular mechanism. Our analysis began with an examination of its amino acid sequence features. A nuclear localization sequence was predicted by NLStradamus tool (Fig. S5A). In addition, the N-terminal region of bHLH121 contains two conserved structural domains: a bHLH domain and a leucine zipper (L-ZIP) domain, which are typically responsible for DNA binding and protein dimerization (Gao & Dubos, 2024; Heim et al., 2003). In contrast, the C-terminal region is intrinsically disordered (Fig. S5B) and highly enriched with potential phosphorylation sites. This structural distinction is supported by a prior study, which employed mass spectrometry to confirm phosphorylation events in the C-terminal region of bHLH121 purified from Fe-deficient plants (Kim et al., 2019).

Subsequently, we conducted a comparative analysis of the physicochemical properties among the full-length protein and two segmented regions, aiming to elucidate how structural features contribute to its functional regulation. Results revealed significant differences between the full-length bHLH121 protein and its N-terminal (N121, residues 1–160) and C-terminal (C121, residues 151–337) segments (Table S1). The full-length protein exhibits a near-neutral isoelectric point (pI, 7.69), moderate instability (instability index: 72.77), and overall hydrophilicity (GRAVY: – 0.964), reflecting a balanced physicochemical profile derived from its two constituent fragments. In contrast, the N-terminal segment shows an acidic pI (6.23), lower instability (instability index: 54.5), and higher aliphatic index (88.38), consistent with a stable, hydrophobic structure that supports DNA binding and protein dimerization via its bHLH-L-ZIP domains. Conversely, the C-terminal segment displays a strongly basic pI (9.01), exceptionally high instability (instability index: 90.31), and the highest hydrophilicity (GRAVY: –1.035), indicating it is a highly flexible, intrinsically disordered region enriched with phosphorylation sites. These properties suggest that the C-terminal fragment may act as a phosphorylation-regulated molecular switch under Fe-limited conditions, where signal-induced modifications could modulate protein activity, stability, or protein-protein interactions, while the N-terminal fragment provides a stable structural platform for DNA binding and transcriptional activation.

### 3.5 Functional analyses of the N- and C-terminal bHLH121 fragments under Fe deficiency

Based on these distinct structural features, we next asked whether the N-terminal or C-terminal fragment was the primary determinant for full-length bHLH121-overexpression-mediated root growth inhibition under Fe deficiency. We generated transgenic plants in WT background overexpressing either the N-terminal 160-amino-acid fragment (N121, containing the bHLH and L-ZIP domains) or the C-terminal 187-amino-acid fragment (C121, spanning the predicted intrinsically disordered region, IDR). Phenotypic analysis was performed using three independent transgenic lines for each of the *N121 ox* and *C121 ox* (Fig. 6A). Notably, *N121 ox* seedlings showed strongly enhanced Fe-deficiency sensitivity, as evidenced by drastically shortened primary roots and severe leaf chlorosis (Fig. 6B). Indeed, N121 overexpression repressed plant growth regardless of Fe sufficiency or deficiency, and exerted a significantly stronger inhibitory effect than full-length bHLH121 overexpression on root growth, seedling fresh weight and chlorophyll content (Figs. 6, S6). Conversely, *C121 ox* plants displayed improved growth phenotype, exhibiting significantly longer primary root than the WT when grown on –Fe medium (Fig. 6B, C). Nevertheless, no obvious difference was observed between WT and *C121 ox* plants under +Fe conditions. Collectively, these results demonstrate that, compared to the full-length bHLH121, the N121 fragment shows dominant nature and exerts an inhibitory effect on plant growth under Fe deficiency.

**Figure 6.**
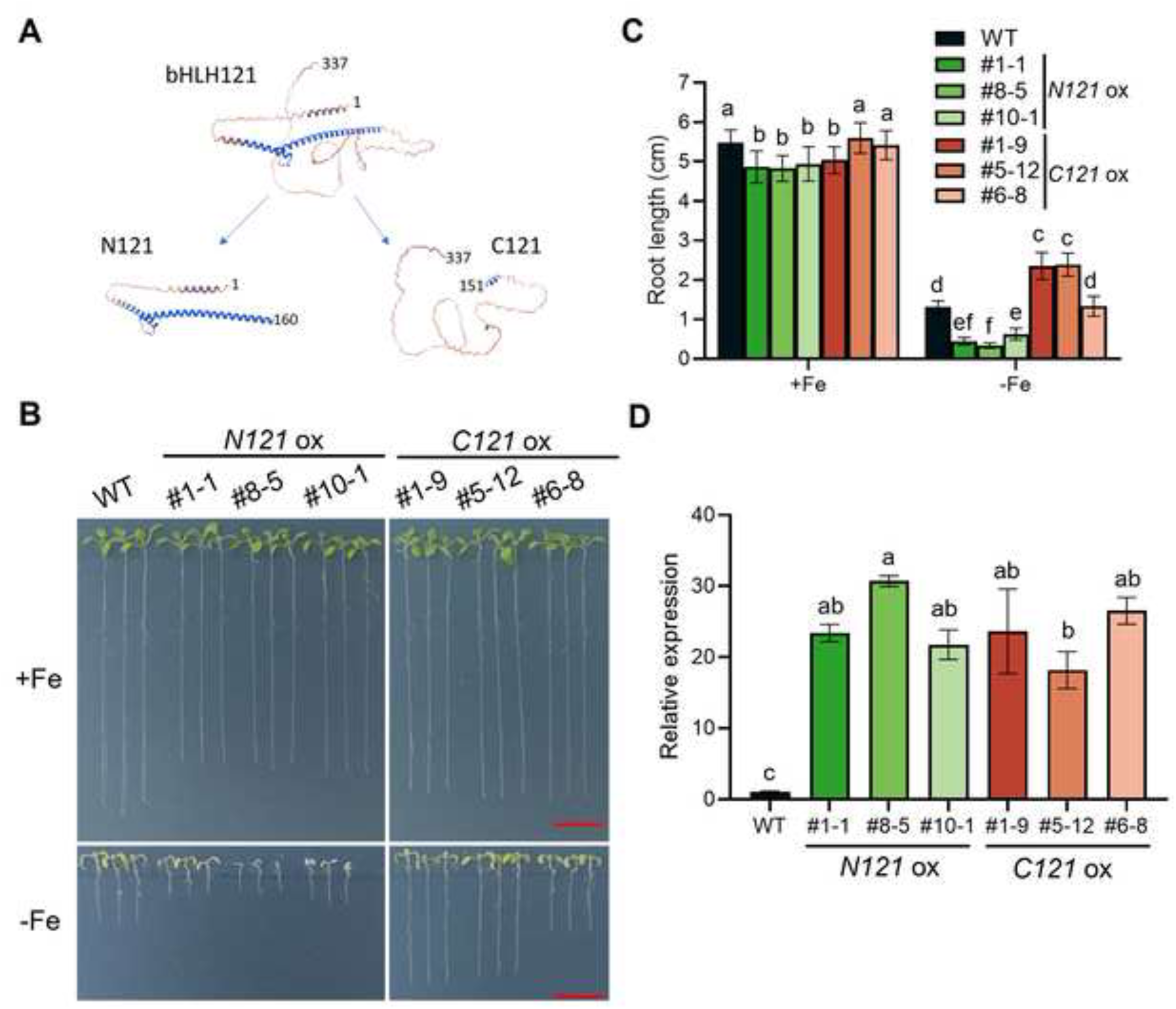
Overexpression of N- and C-terminal *bHLH121* fragment displayed opposing effects on plant tolerance to iron deficiency. (**A**) Schematic diagram of N- and C-terminal domain expression constructs of bHLH121. Based on the structural characteristics of bHLH121 protein, the N-terminal bHLH domain (amino acids 1 to 160) and the C-terminal region containing potential phosphorylation sites (amino acids 151 to 337) were separately introduced into a binary expression vector. (**B**) Phenotypes of 10-day-old seedlings of WT and different transgenic lines grown on +Fe and –Fe medium. Scale bars = 1 cm. (**C**) Root length analysis (n = 30). (**D**) Abundance of *bHLH121* transcripts in WT and transgenic plants. Data are means ± SD. Different letters represent significant differences determined by one-way ANOVA with Tukey’s post hoc test (*P* < 0.05).

### 3.6 Overexpression of bHLH121 and its truncated fragments does not alter plant responses to other mineral nutrient stresses

To determine whether overexpression of *bHLH121* and its truncated fragments alters plant responses to other nutrition stresses, we examined the phenotypes of WT, full-length *bHLH121 ox, N121 ox*, and *C121 ox* lines under low potassium (10 μM K^+^) and phosphorus deficiency (–P). Although a slight increase in low-K resistance was detected in *N121 ox* root growth, no significant phenotype differences were observed among WT, *bHLH121 ox*, *N121 ox*, and *C121 ox* plants grown on –P medium (Fig. S7). Similarly, our results showed that the selected overexpression lines exhibited no significant alterations in root growth in response to deficiencies of the essential metal ions copper (Cu), zinc (Zn), and manganese (Mn) (Fig. S7). Collectively, these data demonstrate that overexpression of bHLH121 and its truncated fragments specifically impacted plant response to Fe deficiency.

### 3.7 Ferric chelate reductase (FCR) activity and expression of Fe uptake genes show significant differences among the tested overexpression lines

To further investigate how overexpression of different bHLH121 variants impacts Fe homeostasis, we analyzed Fe-chelate reductase (FCR) activity using the ferrozine assay (Yi & Guerinot, 1996). Because under Fe conditions, *Arabidopsis* plants typically display an enhanced FCR in the root to mobilize external Fe. Consistent with the previously described Fe deficiency-related phenotypes (Figs. 5 and 6), *N121 ox* plants displayed a substantial decrease in FCR activity compared with the WT, whereas root FCR activity in *C121 ox* plants was comparable to or slightly higher than that in WT and the WT-like *bHLH121 ox* lines, as shown by qualitative and quantitative measurements (Fig. 7A, B). To explore the underlying molecular response in seedlings grown on +Fe and -Fe media for 10 days, we analyzed the transcript levels of *FRO2* and *IRT1*, two key genes responsible for root Fe uptake, via quantitative RT-PCR. As expected, the Fe-deficiency-induced expression of both genes was markedly reduced in *N121 ox* plants (Fig. 7C, D). To further link the observed phenotypes to Fe homeostasis, we measured total Fe concentrations in plants grown under +Fe and 0Fe conditions. Interestingly, total Fe concentrations were comparable among WT, *bHLH121 ox*, and *C121 ox* plants. By contrast, *N121 ox* lines displayed significantly lower Fe content than all other genotypes under both Fe-sufficient and Fe-deficient conditions (Fig. 8), indicating a severely dysfunctional Fe deficiency response. Collectively, these results demonstrate that the N121 substantially represses the Fe deficiency response in *Arabidopsis*.

**Figure 7.**
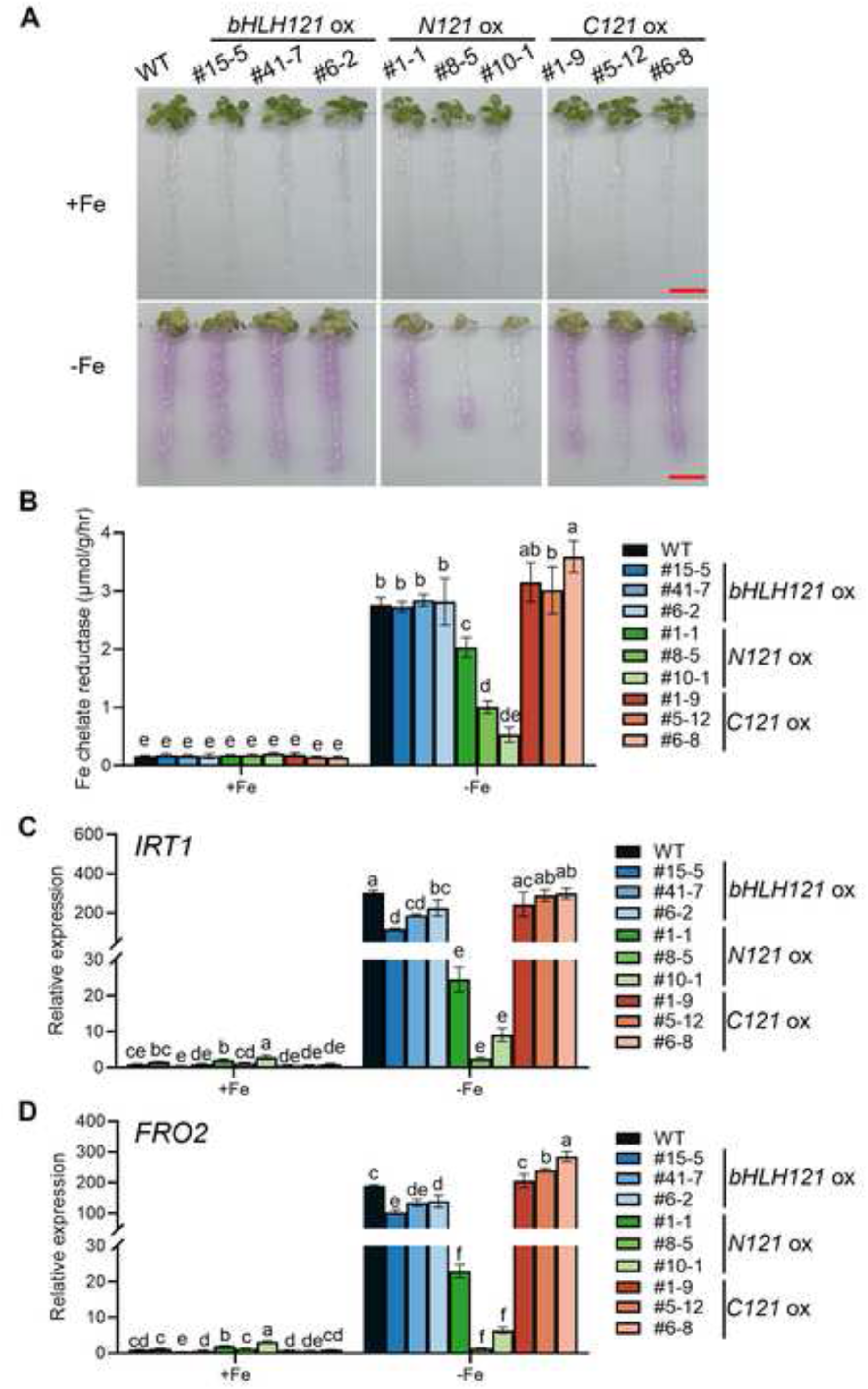
Overexpression of N-terminal *bHLH121* fragment inhibits iron uptake in plants under iron-deficient conditions. (**A**, **B**) Qualitative (**A**) and quantitative (**B**) detection of ferric chelate reductase activity in bHLH121 full-length and segment overexpression lines. Scale bars = 1 cm. Seedlings grown on ES medium for 7 days were transferred to +Fe/-Fe media for 5 days, and root tissues were used for the detection of ferric chelate reductase activity. Data are means ± SD (n = 3). (**C**, **D**) Relative expression of *IRT1* (**C**) and *FRO2* (**D**) in bHLH121 full-length and segmented overexpression lines. Relative expression was determined by RT-qPCR in 10-day-old *Arabidopsis* seedlings grown on +Fe and –Fe medium. Data are means ± SD (n = 3). Different letters represent significant differences determined by one-way ANOVA with Tukey’s post hoc test (*P* < 0.05).

**Figure 8.**
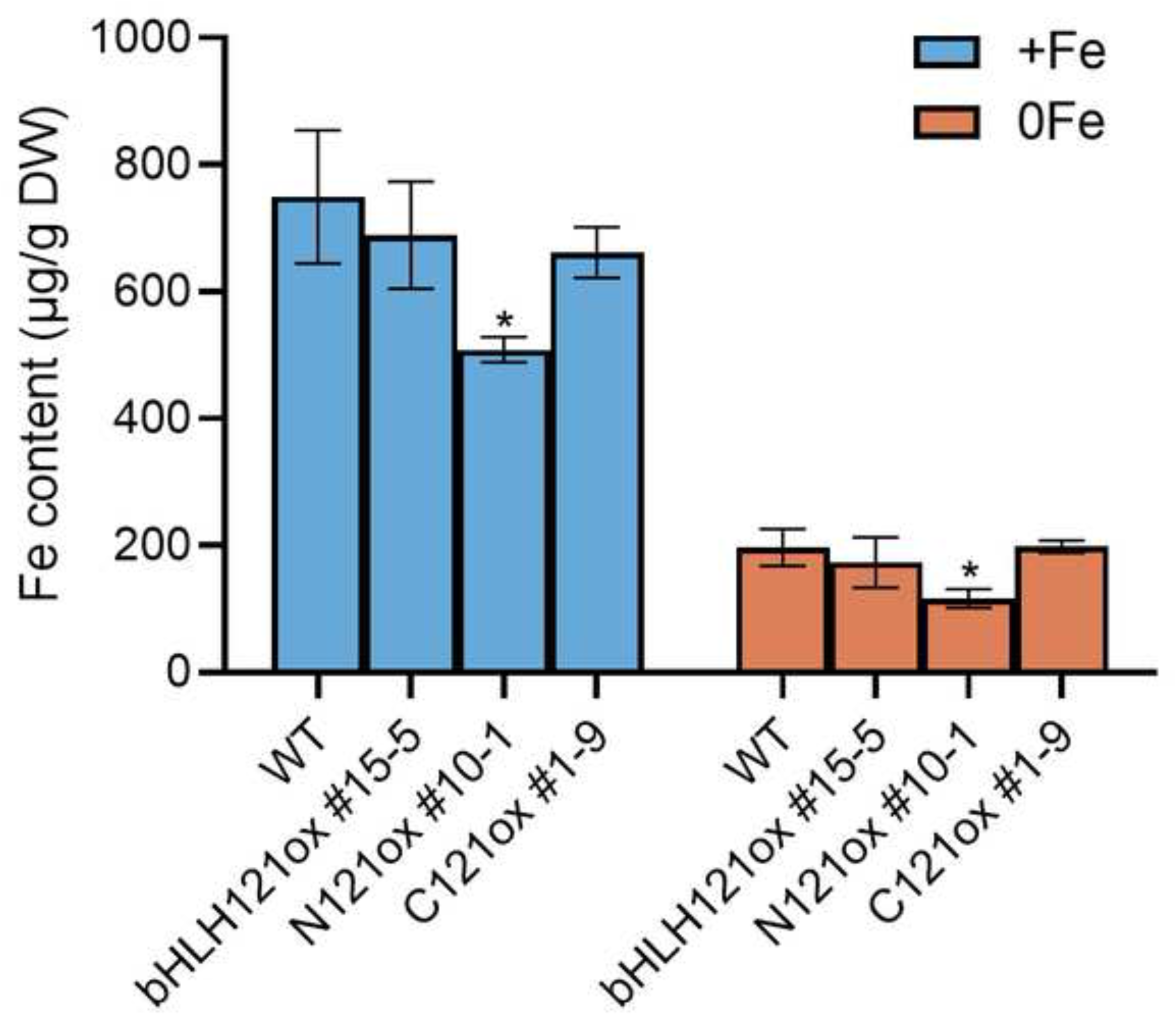
Overexpression of N-terminal *bHLH121* fragment decreases iron content in plants. Fe contents of the bHLH121 full-length and segmented overexpression lines grown on +Fe or -Fe medium for 10 d. Data are means ± SD (n = 3). The unpaired two-tailed Student’s t-test was used to assess the statistical significance of the differences from the WT (\**P* < 0.05).

### 3.8 Transcriptional analysis of several Fe deficiency response regulators in bHLH121 ox, N121 ox, and C121 ox plants

To further determine how bHLH121 variant overexpression mediates the expression of Fe deficiency responsive genes, we performed quantitative real-time PCR analysis using seedlings grown on +Fe and -Fe medium for 10 days. Several representative Fe-deficiency-inducible TFs, including *FIT* and two bHLH Ib subgroup members *bHLH38* and *bHLH39* were selected for expression analysis. All three genes were significantly upregulated in WT plants under Fe-deficient conditions (Fig. 9). Notably, the Fe-deficiency-induced expression of *FIT* in *bHLH121 ox* plants was impaired relative to the WT; in two of the three independent lines, *FIT* transcript levels were significantly lower than in the WT, while the third line exhibited only a modest reduction. Consistent with their severe Fe-deficiency-hypersensitive phenotypes, *FIT* induction was almost completely blocked in all three tested *N121 ox* lines (Fig. 9A). By contrast, the transcript levels of *bHLH38* and *bHLH39* were only slightly modulated in *bHLH121 ox*, *N121 ox* and *C121 ox* plants, although their Fe-deficiency-induced expression was significantly reduced in two of the three tested *N121 ox* lines (Fig. 9B, C). Since *FRO2* and *IRT1* are two key root Fe-uptake genes, whose Fe-deficiency-dependent transcriptional upregulation is directly governed by the FIT-bHLH Ib activation module (Jakoby et al., 2004; Wang et al., 2013; Yuan et al., 2008; Yuan et al., 2005), the impaired induction of *FIT* is likely the primary causal factor underlying the Fe-deficiency-hypersensitive phenotypes observed in the *bHLH121 ox* and *N121 ox* seedlings.

**Figure 9.**
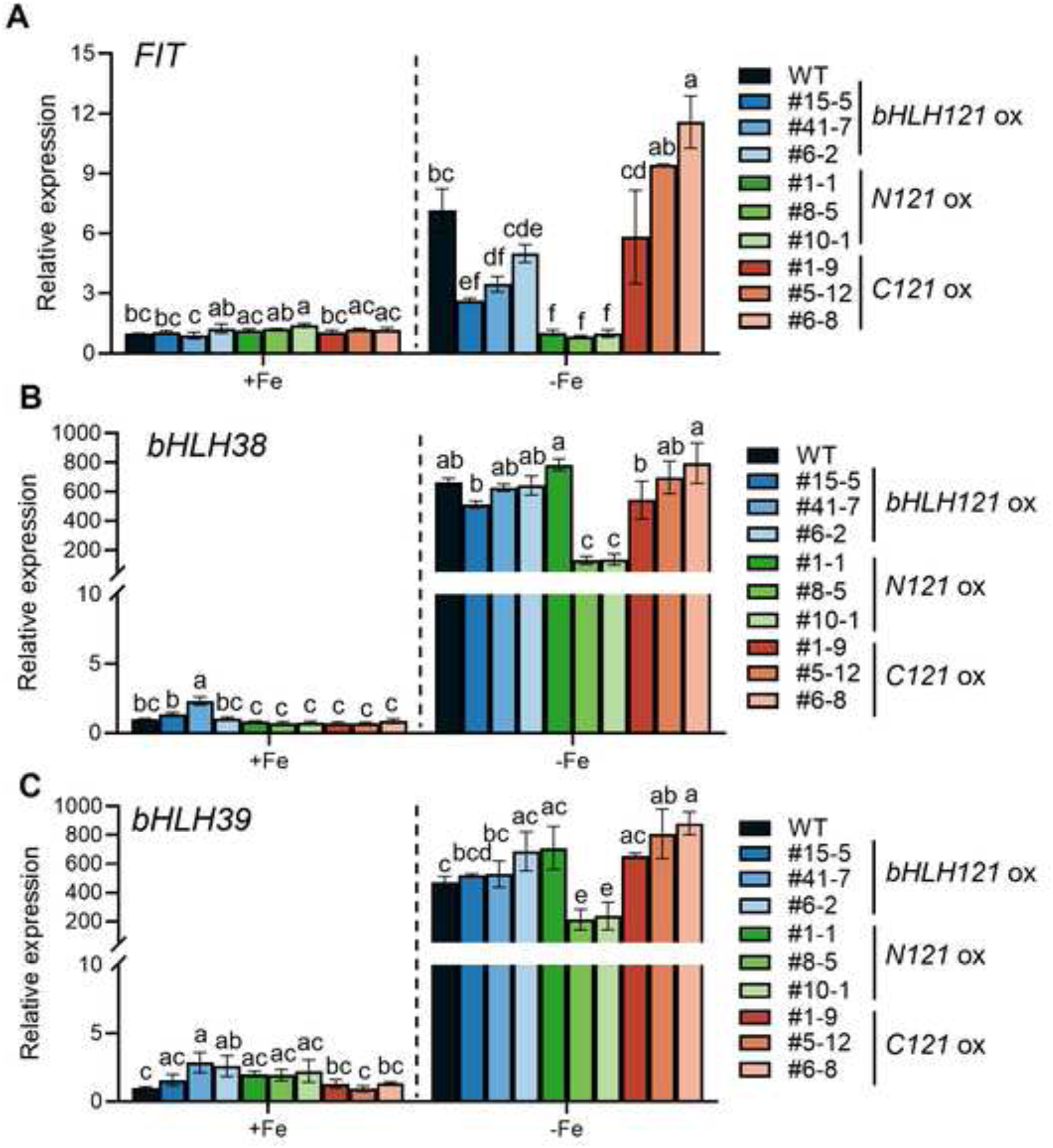
Expression levels of iron-deficiency response genes decreased in *N121 ox* lines under iron-deficient conditions. (**A-C**) Relative expression of *FIT* (**A**)*, bHLH38* (**B**) and *bHLH39* (**C**) in bHLH121 full-length and segmented overexpression lines. Relative expression was determined by RT-qPCR in 10-day-old *Arabidopsis* seedlings grown on +Fe and –Fe medium. Data are means ± SD (n = 3). Different letters represent significant differences determined by one-way ANOVA with Tukey’s post hoc test (*P* < 0.05)

### 3.9 Subcellular localization of N121-GFP and C121-GFP in transgenic plants

To determine the subcellular localization of the ectopically expressed truncated bHLH121 fragment proteins, we fused the green fluorescent protein (GFP) to the C-terminus of N121 and C121, with expression driven by the cauliflower mosaic virus 35S promoter. As an internal control, free GFP signals were detected throughout root cells, localizing to both the cytoplasm and the nucleus (Fig. 10A). Notably, although no canonical nuclear localization signal was predicted in the C121 fragment, GFP fluorescence was observed to preferentially localize to the nucleus in *Arabidopsis* root tip cells expressing C121-GFP, as well the case for the full-length bHLH121-GFP fusion protein (Fig. 10A). Consistent with this observation, previous studies using the endogenous *bHLH121* promoter reported exclusive nuclear localization of bHLH121-GFP (Gao et al., 2020). In sharp contrast, the GFP signal distribution pattern of N121-GFP was distinct from that of bHLH121-GFP, closely mimicking the free GFP signals in both the cytoplasm and the nucleus (Fig. 10A). In addition, we observed a significant reduction in GFP signal intensity for both N121-GFP and C121-GFP but not for bHLH121-GFP, under 0Fe treatment compared with +Fe control (Fig. 10B). This finding suggests that the isolated N- and C-terminal fragments confer protein instability under iron deficiency, potentially via ubiquitin-mediated proteasomal degradation. We note that the diffuse GFP signal observed in several transgenic lines could be attributed to be an artifact caused by high-level ectopic overexpression or properties of the GFP fusion proteins. Thus, we performed quantitative analysis of the nuclear-to-cytoplasmic GFP fluorescence intensity ratio (N/C ratio). The results further confirmed strong nuclear enrichment for the full length bHLH121-GFP fusion protein (N/C ratio > 4) and significant nuclear accumulation for the C121-GFP fusion protein (N/C ratio > 2), whereas N121-GFP (N/C ratio ≈ 1) showed nearly equivalent signal intensities in the nucleus and cytoplasm (Fig. 10C). Taken together, these results support both the N121 and C121 fragments of bHLH121 are capable of nuclear entry, and can interfere with nuclear functions such as gene transcriptional regulation in their respective overexpression lines.

**Figure 10.**
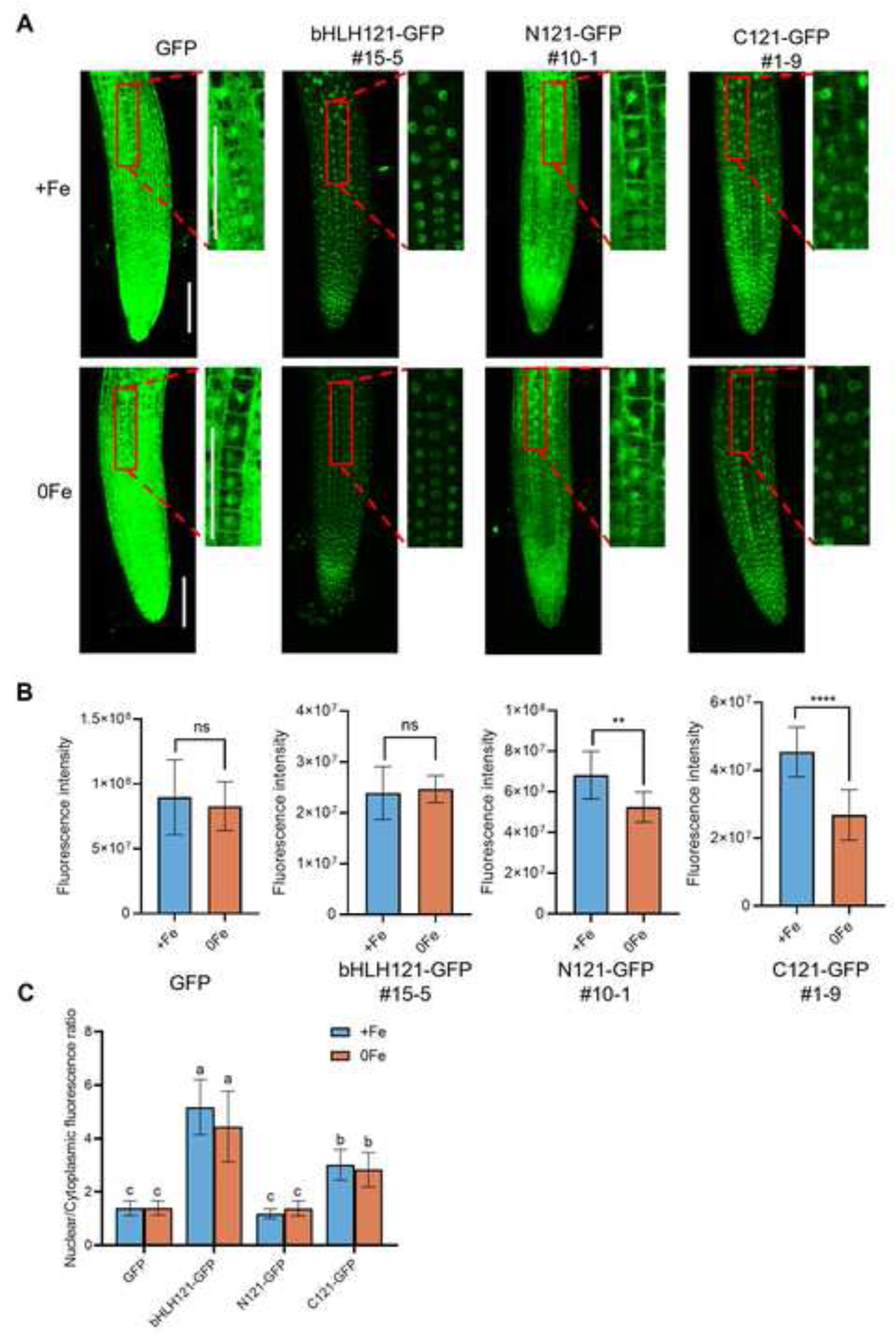
The subcellular localization of bHLH121-GFP, N121-GFP and C121-GFP in transgenic plants. (**A**) GFP fluorescence (green) images are shown. Scale bar = 100 μm. (**B**) GFP fluorescence intensity analysis. Data are means ± SD (n = 10). The unpaired two-tailed Student’s t-test was used to assess the statistical significance of the differences between two groups (ns, non-significant, \*\**P* < 0.01, and \*\*\*\**P* < 0.0001). (**C**) Nuclear/cytoplasmic fluorescence ratio. Different letters represent significant differences determined by one-way ANOVA with Tukey’s post hoc test (*P* < 0.05). Consistent results were obtained across three independent biological replicates.

### 3.10 N-terminal bHLH121 protein can compete with full-length bHLH121 for FIT promoter binding in vitro

Given that bHLH121 has been reported to directly bind the *FIT* promoter and is required for the activation of *FIT* transcription by bHLH IVc proteins (Lei et al., 2020), while two other independent studies failed to detect such in planta DNA-protein interaction (Gao et al., 2020; Kim et al., 2019), we hypothesized that the ectopically expressed N121 antagonistically suppressed this endogenous bHLH121-mediated activation process in *N121 ox* plants. To test this hypothesis, we performed an electrophoretic mobility shift assay (EMSA) using purified bacterially expressed His-TF-tagged full-length bHLH121, N121 and C121 recombinant proteins (Fig. S8). EMSA results showed that both full-length bHLH121 and the N121 fragment could effectively bind the *FIT* promoter-derived probes (Fig. 11A). In sharp contrast, the C121 fragment completely failed to associate with the *FIT* promoter probes (Fig. 11A, B). When N121 was co-incubated with full-length bHLH121 and the *FIT* promoter probe, an increased abundance of the protein-DNA complex was observed, further confirming that both N121 and bHLH121 possesses intrinsic DNA-binding ability. In contrast, the addition of C121 did not significantly alter the intensity of the bHLH121-probe complex (Fig. 11B). Notably, at a higher protein concentration (2×), C121 also slightly reduced the abundance of full-length bHLH121-probe complex (Fig. 11B), suggesting a possible non-specific or concentration-dependent interference effect, although this effect was far less pronounced than that observed for N121. Together, these data demonstrate that the N121 fragment can effectively compete with full-length bHLH121 for binding to the *FIT* promoter *in vitro*.

**Figure 11.**
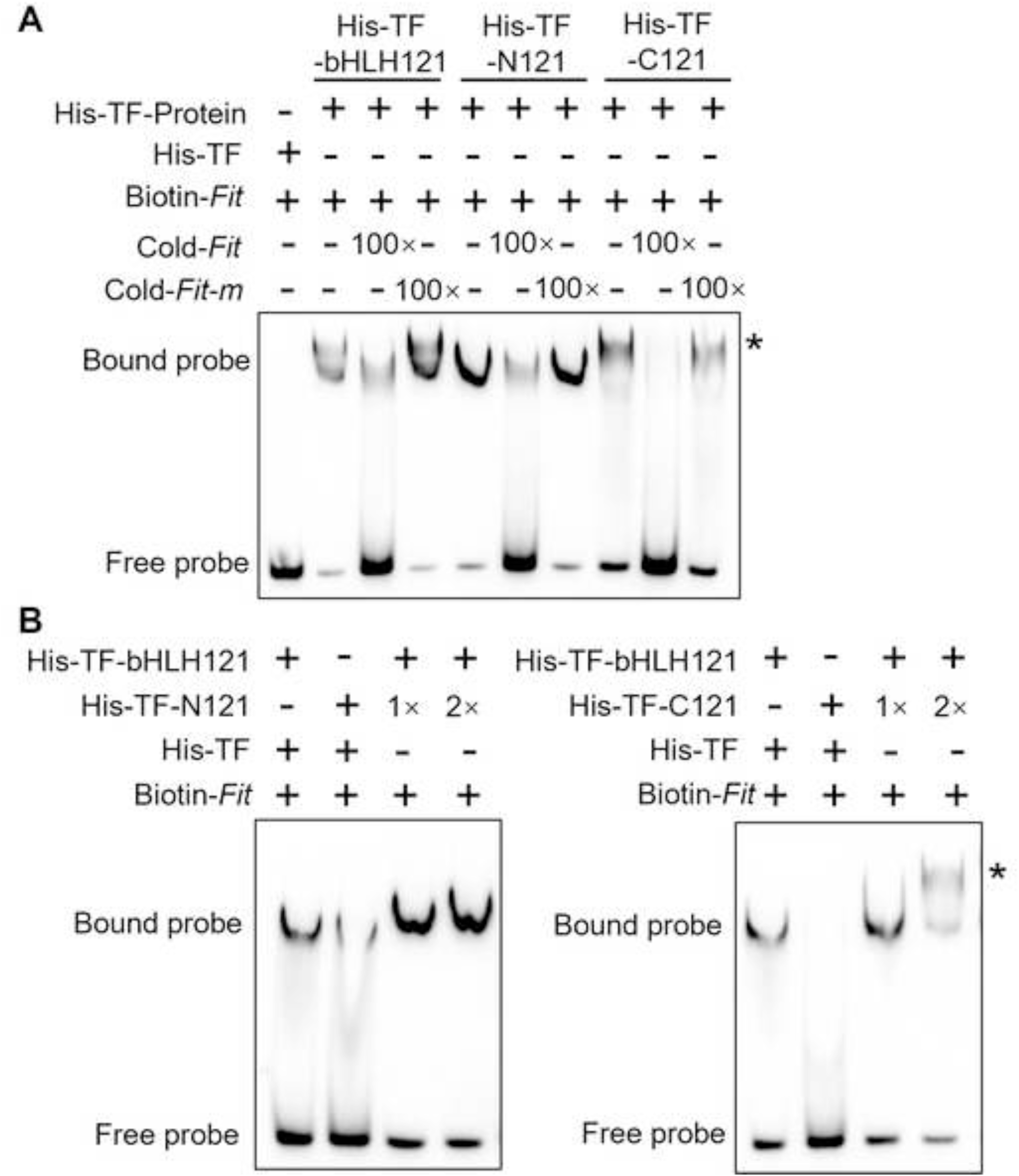
The N-terminal bHLH121 protein binds the *FIT* promoter *in vitro*. **(A)** EMSA results showing that the bHLH121 and N121 protein directly bind the *FIT* promoter, whereas the C121 protein cannot. **(B)** EMSA results showing that co-incubation with the N121 protein enhanced binding to the *FIT* promoter, whereas the C121 protein cannot. Biotin-Probe, biotin-labeled probe; Cold-Probe, unlabeled probe; Cold-Probe-m, unlabeled probe with mutated E-box. Biotin probe incubated with His-TF derived from empty vector served as the negative control. Asterisks indicate non-specific bands for the C121 protein.

## 4. Discussion

The maintenance of cellular Fe homeostasis represents a core physiological process that enables plants to survival and reproductive success in dynamically fluctuating environments. As an essential micronutrient, Fe plays irreplaceable functional roles in a wide spectrum of fundamental biological activities such as electron transport chains, cellular energy metabolism, and hormone synthesis. However, its inherent dual nature—acting both as an indispensable nutrient and a potent trigger of reactive oxygen species (ROS) toxicity via the Fenton reaction, has driven land plants to evolve an exquisitely sophisticated, multi-layered, and highly integrated Fe regulatory network. This study, employing a unique experimental strategy of ectopically expressing distinct truncated variants of the transcription factor bHLH121/URI, to systematically investigate both its evolutionarily conserved function across multiple plant species and the distinct regulatory roles mediated by its independent N-terminal and C-terminal segments. By integrating multi-layered phenotypical, genetic, physiological, and molecular analyses, the present work not only uncovers the conserved function of bHLH121 in several dicotyledonous species but also demonstrates that overexpression of its N-terminal fragment can strongly attenuate Fe deficiency response in *Arabidopsis thaliana*.

The central, multifaceted role of the bHLH transcription factor family in regulating plant Fe homeostasis is well-established, forming an evolutionarily conserved, complex hierarchical regulatory cascade network that extends from master regulator FIT and clade IVc bHLHs (e.g., bHLH105/ILR3, bHLH115) in *Arabidopsis* to the functionally orthologous IRO2 transcription factor in gramineous crops (Liang, 2022). Recent independent studies have placed bHLH121/URI at the functional core of this Fe regulatory network (Gao et al., 2024; Gao et al., 2020; Kim et al., 2019; Lei et al., 2020). Distinct from other well-characterized bHLH regulators in this pathway, bHLH121 exhibits two highly unusual regulatory mode. First, unlike the vast majority Fe-responsive genes, its transcripts are constitutively expressed across all major plant tissues irrespective of Fe availability, suggesting that its functional regulation occurs primarily at the post-transcriptional or post-translational level, rather than through transcriptional induction under Fe deficiency. Second, its tissue-level distribution and subcellular localization are dynamically modulated by cellular Fe status, with pronounced enrichment of bHLH121 in the nucleus of root cortex and rhizodermis cells under Fe deficiency (Gao et al., 2020; Lei et al., 2020), a process that has been hypothesized to be mediated by targeted protein phosphorylation (Kim et al., 2019). Importantly, beyond its well-characterized and central role in Fe homeostasis, bHLH121 has recently been functionally implicated in additional abiotic stress response pathways. A recent study in peanut demonstrated that AhbHLH121 positively regulates salt tolerance by directly activating the expression of ROS-scavenging enzyme genes (Zhao et al., 2024). In the present study, a large set of putative bHLH121 homologs across the plant lineage were identified via comprehensive sequence alignment. Our subsequent functional validation assays further support the occurrence of partial functional divergence, as the four selected bHLH121 homologs we tested exhibited quantitatively distinct capacities to rescue the characteristic Fe-deficient phenotype of the *Arabidopsis bhlh121* loss-of-function mutant (Figs. 2 and 4). These evolutionarily divergent roles of bHLH121 homologs from different plant species, in the context of Fe homeostasis and/or other stress responses clearly warrant dedicated systematic investigation in future work. Such efforts will not only provide novel mechanistic insights into the evolutionary diversification of this critical transcription factor clade, but also greatly facilitate the translational application of these homologous genes in next-generation crop improvement for enhanced nutrient use efficiency and multiple stress tolerance.

The present study demonstrates that representative *bHLH121-GFP* complementation lines, such as #34-3, which accumulate *bHLH121-GFP* fusion transcripts at levels drastically exceeding the endogenous *bHLH121* transcript abundance in wild-type (WT) plants, exhibit significantly elevated ferric chelate reductase (FCR) activity under Fe-deficient conditions compared with the WT control (Fig. 1). In sharp contrast, the 35S promoter-driven full-length *bHLH121* overexpression lines display FCR activity that is either comparable to or even lower than that of the WT under identical -Fe treatment conditions (Fig.7). Notably, both sets of transgenic lines effectively overexpress the *bHLH121* coding sequence at the transcript level, yet they produce completely inconsistent physiological outcomes at the functional level. We reason that the line #34-3 expresses the bHLH121-GFP fusion protein under the control of its native endogenous promoter, whereas the 35S-driven overexpression lines accumulate this recombinant protein under the direction of the constitutive CaMV 35S promoter. This fundamental difference in transcriptional regulatory elements could drastically alter the spatiotemporal expression pattern of *bHLH121* across different root tissues and cell types, ultimately resulting in the markedly distinct FCR activity profiles observed under Fe-deficient conditions. We therefore emphasize that the overexpression phenotypes of key regulatory transcription factors such as bHLH121 are highly context-dependent, and that native promoter-driven transgene expression far better recapitulates the authentic spatial and temporal dynamics of endogenous gene regulation. In addition. we note that the *in vivo* protein accumulation levels of bHLH121 are not necessarily proportional to its steady-state transcript abundance, and that the observed GFP fluorescence signal alone cannot formally rule out partial proteolytic cleavage and truncation of the full-length fusion protein. The potential uncharacterized differences in total protein abundance, subcellular partitioning, and protein stability among these distinct transgenic lines clearly remain to be systematically dissected in future dedicated investigations.

Overexpression of the N-terminal fragment of *bHLH121* (hereafter designated *N121 ox*) renders transgenic plants significantly more hypersensitive to Fe limitation, exhibiting far more severe root growth inhibition and leaf chlorosis than WT. The potent inhibitory function of the N121 is likely mediated through two mutually non-exclusive mechanistic routes. First, the ectopically overproduced N121 protein acts by competitively sequestering the previously well-documented interaction partners of full-length bHLH121, such as the clade IVc bHLH transcription factors. This mode of action is highly analogous to a previously reported mechanism, in which overexpression of full-length bHLH121 was shown to suppress the Fe-deficiency response by forming non-functional bHLH121 homodimers that interfere with the assembly of the physiologically critical bHLH121-IVc bHLH functional heterodimers (Lei et al., 2020). Second, the N-terminal region of bHLH121 itself constitutes the key functional domain responsible for target gene recognition and DNA binding, as the N121 fragment completely harbors the canonical bHLH DNA-binding domain that defines this transcription factor family. Overexpression of the N-terminal fragment N121 is proposed to exert its inhibitory effect by directly competing with endogenous full-length bHLH121 for overlapping cis-regulatory DNA binding sites at its target gene promoters. Notably, we have confirmed that purified N121 protein can specifically associate with the conserved E-box motifs within the *FIT* promoter sequence *in vitro* via electrophoretic mobility shift assay (Fig. 11), but the detailed in planta mechanisms underlying this competitive regulatory action remain to be fully elucidated. Moreover, complementary *in vivo* experiments including quantitative ChIP-qPCR and targeted promoter-luciferase transactivation assays will be strictly required to formally validate this competitive dominant-negative working model. To further rigorously test the genetic specificity of the N121-mediated inhibitory effect, it will be functionally essential to express the *N121* transgene specifically in the homozygous *bhlh121* null mutant background, and quantitatively determine whether the characteristic Fe-hypersensitive inhibitory phenotype of N121 overexpression lines is completely attenuated in the absence of endogenous full-length bHLH121. On the other side, transgenic lines overexpressing the isolated C-terminal fragment of *bHLH121* (designated *C121 ox*) exhibited significantly enhanced primary root growth compared with wild-type (WT) plants under consistent Fe-deficient growth conditions (Fig. 6). The C-terminal region of bHLH121 is predicted to be an intrinsically disordered polypeptide with a strongly basic theoretical pI of 9.01, a biochemical property that can promote non-specific electrostatic interactions with negatively charged DNA backbones or pre-formed protein-DNA complexes at elevated local protein concentrations. The observed reduction in the abundance of the full-length bHLH121-DNA probe complex in our EMSA assays at high C121 concentrations (Fig. 11) most likely reflects such non-specific biophysical effects, including non-specific protein aggregation or charge-based interference, rather than sequence-specific competitive DNA binding. Consistent with this interpretation, our experimental data confirm that the C121-GFP fusion protein localizes predominantly to the plant cell nucleus (Fig. 10), while purified C121 protein alone shows no detectable direct binding to the *FIT* promoter sequence *in vitro* (Fig. 11). Despite these observations, the precise in planta physiological role of the C121 fragment during Fe homeostasis regulation remains currently unresolved.

A central unresolved question emerging from this work is how the N-terminal and C-terminal regions of bHLH121, as two functionally distinct modular segments of the same full-length transcription factor, coordinately orchestrate its overall regulatory function in wild-type plants. Given the well-established paradigm that bHLH121 executes its core regulatory roles in Fe homeostasis through direct heterodimerization with clade IVc bHLH transcription factors (Kim et al., 2019; Gao et al., 2020; Lei et al., 2020), and that site-specific phosphorylation events located within its C-terminal region could exert a strong positive modulatory effect on its transcriptional activity (Kim et al., 2019), we propose a working model in which the N-terminal and C-terminal domains of bHLH121 physically interact with completely distinct sets of upstream regulatory proteins. Under Fe-sufficient conditions, bHLH121 is maintained in a low-activity state via physical association with currently uncharacterized negative regulatory proteins, and consequently exhibits only minimal transcriptional activity toward the Fe uptake gene battery. Nevertheless, even under this repressive condition, bHLH121 remains strictly required for other essential physiological processes such as intercellular Fe distribution and intracellular Fe compartmentalization, a functional role that is strongly supported by its previously documented positive regulatory function in controlling the expression of *Ferritin* family genes (Gao et al., 2020). Under Fe-deficient conditions, upstream cellular signaling cascades, most notably targeted phosphorylation events, are proposed to post-translationally modify the C-terminal region of bHLH121. This covalent modification allosterically disrupts the repressive protein-protein interaction, thereby liberating the N-terminal domain of bHLH121 to engage in productive, high-affinity heterodimerization with its clade IVc bHLH partner proteins. Under Fe-sufficient conditions, the inhibitory action of these currently uncharacterized negative regulatory factors is functionally dominant, and is proposed to be mediated through two mutually compatible mechanisms: (1) physically blocking the effective protein-protein interaction between bHLH121 and IVc bHLH transcription factors, or directly interfering with the subsequent recruitment of the basal transcriptional machinery; and (2) sequestering the majority of cellular bHLH121 protein in the cytosol, or partitioning it into catalytically inactive higher-order protein complexes. As a direct physiological consequence, the entire suite of downstream Fe acquisition genes remains tightly transcriptionally repressed, preventing unnecessary high-affinity Fe uptake and the consequent risk of cellular Fe overload and oxidative toxicity. Upon specific perception of Fe deficiency, upstream stress signaling pathways trigger targeted post-translational modification of bHLH121, most likely via site-specific phosphorylation within its C-terminal tail. This modification event operates as a critical “molecular switch” that dynamically shifts the binding equilibrium of bHLH121 away from its negative regulators and toward its activating IVc bHLH partners. This regulatory switch can potentially relieve the inhibitory protein association, expose the otherwise masked protein-protein interaction and transcriptional activation interfaces, or alter the subcellular localization dynamics of bHLH121 to promote its efficient nuclear import. Once inside the nucleus, the post-translationally modified bHLH121 protein efficiently assembles into functional heterodimeric complexes with IVc bHLH proteins, which themselves accumulate to high abundance under Fe-deficient conditions due to the well-documented attenuation of their BTS E3 ligase-mediated proteasomal degradation pathway. This transcriptionally active bHLH121-IVc bHLH complex subsequently binds to the E-box cis-regulatory motifs in the promoter regions of core Fe regulatory genes such as *FIT* and *bHLH38/39*, resulting in the robust and coordinated transcriptional activation of the plant’s high-affinity Fe acquisition response program.

Despite the substantial new insights provided by the present study, the precise multifaceted molecular mechanisms that underpin the full regulatory function of bHLH121 in plant Fe homeostasis remain to be fully systematically elucidated. A series of high-priority, interconnected key scientific questions clearly warrant dedicated future investigation. First, integrative structural biology approaches, including both X-ray crystallography and single-particle cryo-electron microscopy, combined with rigorous quantitative biochemical assays, will be strictly required to precisely resolve the atomic-level interaction interfaces and the underlying structural basis that governs the physical associations of full-length bHLH121, as well as its isolated N-terminal and C-terminal modular segments, with clade IVc bHLH transcription factors (Lei et al., 2020;

Gao et al., 2024), the basal general transcription machinery, and the currently unidentified putative co-repressor and co-activator protein partners. Second, it will be critically important to systematically map all specific, functionally critical post-translational modification sites on bHLH121, including phosphorylation, ubiquitination, and other understudied regulatory marks, under both Fe-sufficient and Fe-deficient physiological conditions, and to unambiguously identify the full suite of upstream regulatory enzymes involved, including the cognate kinases and phosphatases, as well as the corresponding E3 ubiquitin ligases and deubiquitinases, and to fully dissect their respective upstream signaling regulatory mechanisms (Gao & Dubos, 2024; Li et al., 2022; Zhang et al., 2023). Third, the development of advanced *in vivo* live-cell imaging and genetically encoded FRET biosensor tools (Martin et al., 2021) will enable researchers to monitor, with high spatiotemporal resolution, the real-time protein conformational dynamics, dynamic subcellular localization shuttling, differential protein stability, and transient protein-protein interaction kinetics of bHLH121 specifically in key Fe-responsive tissues such as root tips, directly revealing its *in vivo* real-time response kinetics to dynamic fluctuations in external Fe availability. Fourth, it remains an open and high-impact question to systematically investigate whether and how the bHLH121 regulatory node functions as a central signal integrator to coordinate multiple other endogenous and environmental signals, including the availability of other essential macronutrients such as phosphorus (Xue et al., 2023), and micronutrients such as zinc and copper, as well as diverse hormone signaling pathways, and to rigorously test the hypothesis that the functional equilibrium between its N-terminal and C-terminal domains acts as the critical molecular convergence point for this multi-signal integration process. Fifth, there is also a pressing need to systematically explore the conserved and diversified biological functions and underlying regulatory mechanisms of bHLH121 homologs in agronomically important gramineous cereal crops such as wheat and maize, as well as in evolutionarily divergent plant species that employ fundamentally distinct Fe acquisition strategies (Pan et al., 2024; Gao et al., 2026; Meng et al., 2026), and to comprehensively assess the translational biotechnological potential of these bHLH121 homologs as broad-spectrum high-value molecular targets for the genetic improvement of crop Fe nutritional quality and Fe use efficiency in low-Fe agricultural soils.

## 5. Conclusion

The present study comprehensively investigated the functional conservation and distinct physiological roles of different structural modular forms of the bHLH121 transcription factor, a central regulator of plant Fe homeostasis. Our comparative functional assays demonstrate that the core Fe regulatory function of bHLH121 is evolutionarily conserved across multiple representative dicotyledonous plant species. We further provide multiple lines of *in vitro* and *in planta* experimental evidence demonstrating that the isolated N-terminal fragment of bHLH121 (designated N121) exerts an inhibitory effect on the transcriptional activation of the Fe deficiency response program, strongly suppressing the rapid induction of key Fe homeostasis regulatory genes including *FIT*, and consequently significantly impairing physiological Fe accumulation in planta. Although we have confirmed that N121 protein can directly associate with the *FIT* promoter sequence *in vitro* via electrophoretic mobility shift assays, the underlying molecular mechanism of this inhibitory effect is most likely mediated through competitive DNA binding. Nevertheless, alternative mechanistic possibilities, such as the effective sequestration of endogenous clade IVc bHLH protein interaction partners by N121, cannot be formally excluded at this stage. Collectively, the findings presented in this work clarify the previously uncharacterized functional significance of the N-terminal domain of bHLH121, and provide a solid foundational experimental framework to guide future investigations into the multilayered regulatory complexity of this critical plant Fe homeostasis transcription factor.

## CRediT authorship contribution statement

**Peijun Zhou**: Writing – original draft, Investigation. **Chuanfa Liu**: Writing – original draft, Writing – review & editing, Funding acquisition, Conceptualization. **Yilin Pan**: Investigation. **Yuchen Fei**: Investigation**. Renfang Shen**: Resources. **Ruonan Wang:** Resources, Funding acquisition, Conceptualization. **Ping Lan**: Writing – review & editing, Supervision, Resources, Funding acquisition, Conceptualization. All authors read and approved the final manuscript.

## Acknowledgements

We are grateful to Dr. Fei Gao from Hunan Agricultural University for critical comments. We thank Ms Rong Huang from the Analysis Center for technical assistance. Thanks are given to Misses Xinran Du and Bingyan Liu from the laboratory for their critical comments and useful discussion. The present study was funded by the National Natural Science Foundation of China (grant numbers: 32300274, 32070279), Natural Science Foundation of Jiangsu Province (grant number: BK20221560), and Project of Priority and Key Areas, Institute of Soil Science, Chinese Academy of Sciences (ISSASIP2222, ISSASIP2206).

## Appendix A. Supporting information

**Figure S1** Generation of *bhlh121* loss-of-function mutants via the CRISPR/Cas9 genome editing system

**Figure S2** Phylogenetic and conserved sequence identity analyses of putative bHLH121 homologs identified from ten dicotyledonous plant species

**Figure S3** Functional complementation effects of ectopically expressed bHLH121 homologs in the *bhlh121-7* loss-of-function mutant background

**Figure S4** A statistically significant negative correlation between the relative transcript abundance of *bHLH121* and primary root elongation length under iron deprivation conditions

**Figure S5** Full-length amino acid sequence characterization and intrinsically disordered region prediction analysis of the bHLH121 transcription factor

**Figure S6** Constitutive overexpression of the isolated N-terminal bHLH121 fragment confers severe plant growth inhibition, independent of cellular iron sufficiency or deficiency status

**Figure S7** Transgenic lines overexpressing the isolated N-terminal fragment of bHLH121 are phenotypically insensitive to a broad range of non-Fe macronutrient and micronutrient deficiency conditions

**Figure S8** SDS-PAGE purity and integrity validation of recombinant His-TF-tagged fusion proteins used for electrophoretic mobility shift assays

**Table S1** Systematic physicochemical property characterization of full-length bHLH121 and its isolated N- and C-terminal truncated fragments

**Table S2** Primers used in this study

## Data availability

All data supporting the findings of this study are available within the paper and within its supplementary materials published online.

