## supplemental files for "The conserved function of bHLH121 and overexpression of N-terminal bHLH121 fragment suppresses iron deficiency response in *Arabidopsis*"

### Slide 1
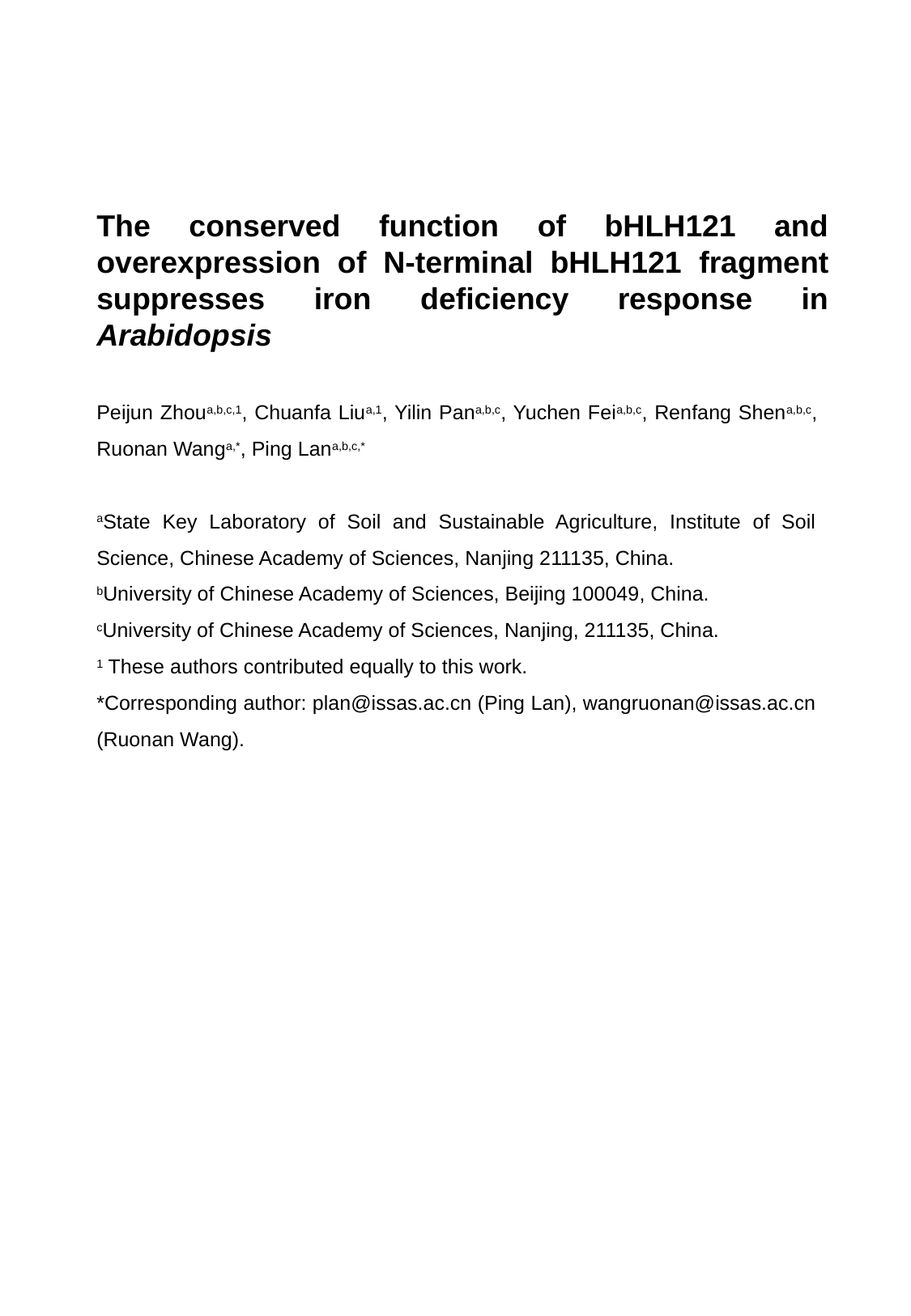

The conserved function of bHLH121 and overexpression of N-terminal bHLH121 fragment suppresses iron deficiency response in Arabidopsis
Peijun Zhou^a,b,c,1^, Chuanfa Liu^a,1^, Yilin Pan^a,b,c^, Yuchen Fei^a,b,c^, Renfang Shen^a,b,c^, Ruonan Wang^a,*^, Ping Lan^a,b,c,*^
^a^State Key Laboratory of Soil and Sustainable Agriculture, Institute of Soil Science, Chinese Academy of Sciences, Nanjing 211135, China.
^b^University of Chinese Academy of Sciences, Beijing 100049, China.
^c^University of Chinese Academy of Sciences, Nanjing, 211135, China.
^1^ These authors contributed equally to this work.

### Slide 2
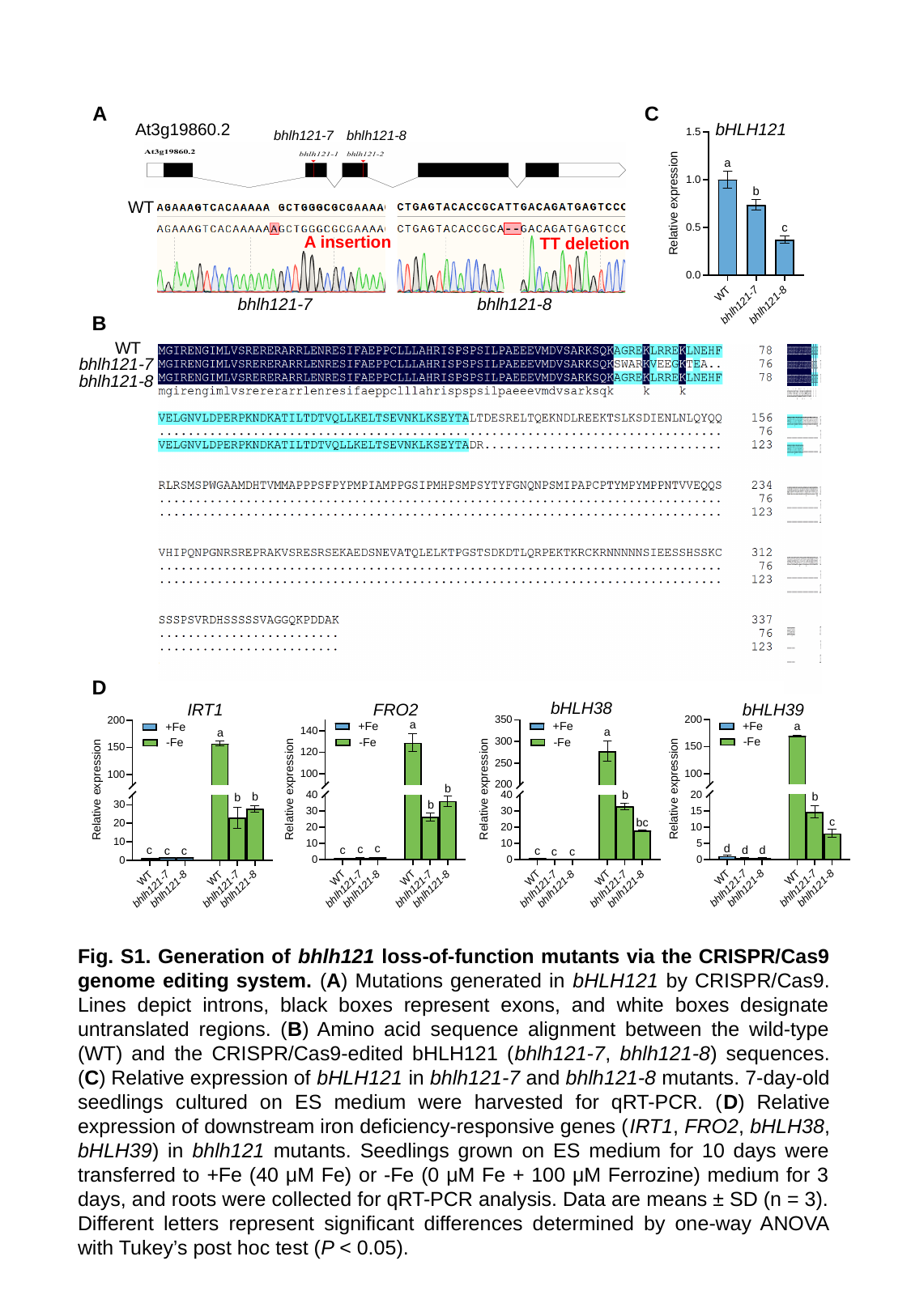

A
C
At3g19860.2
bhlh121-8
bhlh121-7
WT
A insertion
TT deletion
bhlh121-8
bhlh121-7
B
WT
bhlh121-7
bhlh121-8
bHLH121
D
bHLH38
IRT1
FRO2
bHLH39
Fig. S1. Generation of bhlh121 loss-of-function mutants via the CRISPR/Cas9 genome editing system. (A) Mutations generated in bHLH121 by CRISPR/Cas9. Lines depict introns, black boxes represent exons, and white boxes designate untranslated regions. (B) Amino acid sequence alignment between the wild-type (WT) and the CRISPR/Cas9-edited bHLH121 (bhlh121-7, bhlh121-8) sequences. (C) Relative expression of bHLH121 in bhlh121-7 and bhlh121-8 mutants. 7-day-old seedlings cultured on ES medium were harvested for qRT-PCR. (D) Relative expression of downstream iron deficiency-responsive genes (IRT1, FRO2, bHLH38, bHLH39) in bhlh121 mutants. Seedlings grown on ES medium for 10 days were transferred to +Fe (40 μM Fe) or -Fe (0 μM Fe + 100 μM Ferrozine) medium for 3 days, and roots were collected for qRT-PCR analysis. Data are means ± SD (n = 3). Different letters represent significant differences determined by one-way ANOVA with Tukey’s post hoc test (P < 0.05).

### Slide 3
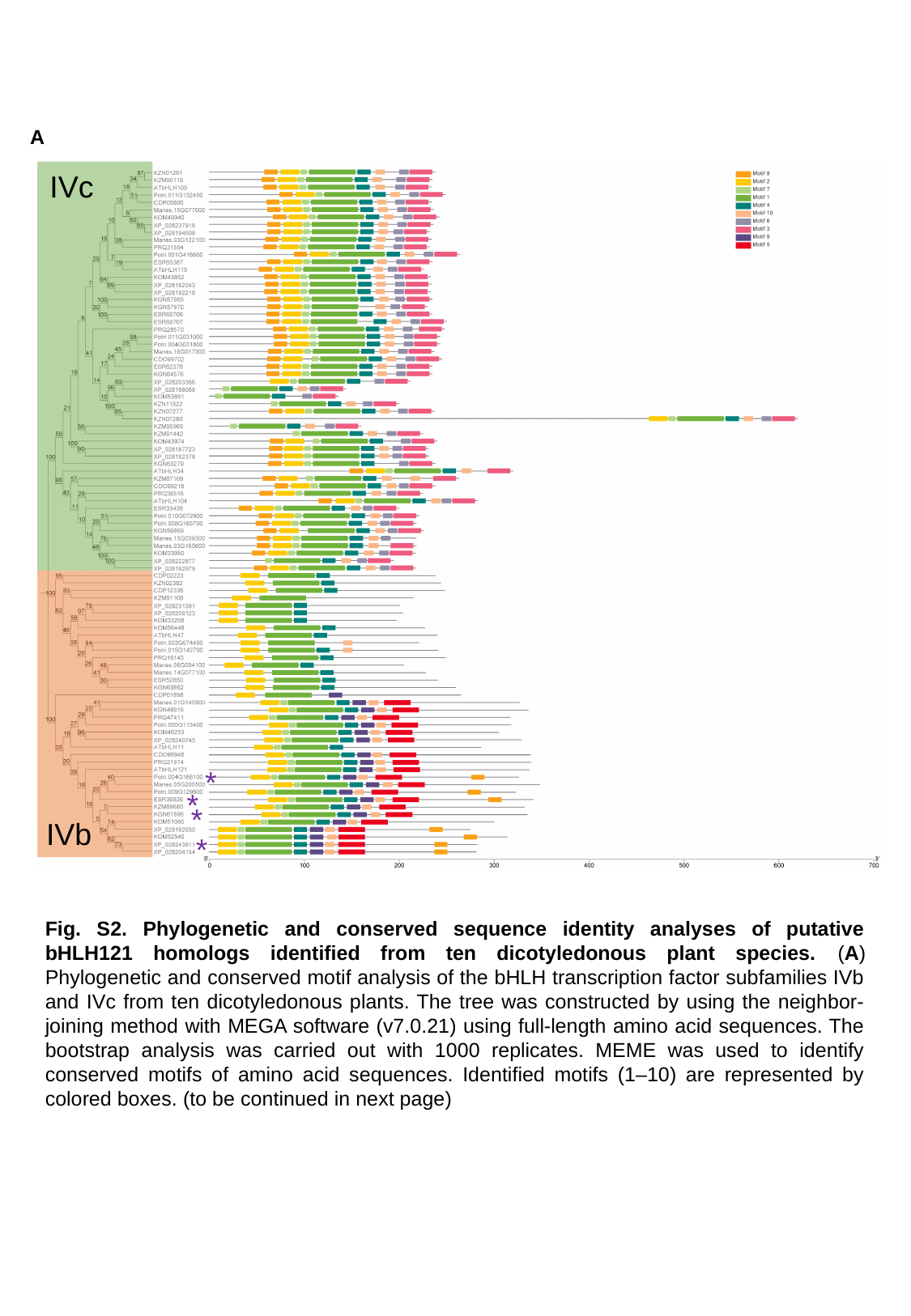

A
IVc
*
*
*
IVb
*
Fig. S2. Phylogenetic and conserved sequence identity analyses of putative bHLH121 homologs identified from ten dicotyledonous plant species. (A) Phylogenetic and conserved motif analysis of the bHLH transcription factor subfamilies IVb and IVc from ten dicotyledonous plants. The tree was constructed by using the neighbor-joining method with MEGA software (v7.0.21) using full-length amino acid sequences. The bootstrap analysis was carried out with 1000 replicates. MEME was used to identify conserved motifs of amino acid sequences. Identified motifs (1–10) are represented by colored boxes. (to be continued in next page)

### Slide 4
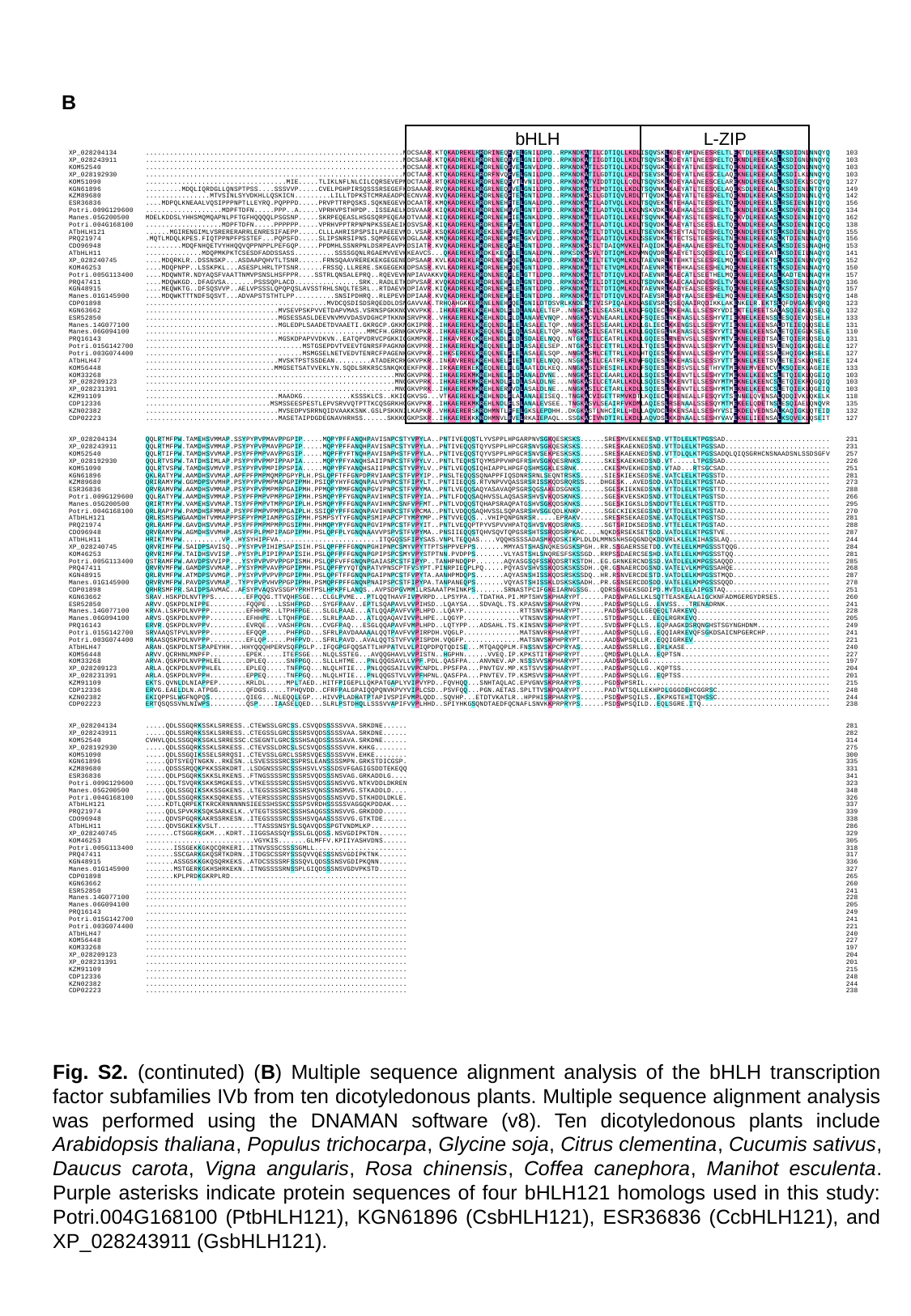

B
bHLH
L-ZIP
Fig. S2. (continuted) (B) Multiple sequence alignment analysis of the bHLH transcription factor subfamilies IVb from ten dicotyledonous plants. Multiple sequence alignment analysis was performed using the DNAMAN software (v8). Ten dicotyledonous plants include Arabidopsis thaliana, Populus trichocarpa, Glycine soja, Citrus clementina, Cucumis sativus, Daucus carota, Vigna angularis, Rosa chinensis, Coffea canephora, Manihot esculenta. Purple asterisks indicate protein sequences of four bHLH121 homologs used in this study: Potri.004G168100 (PtbHLH121), KGN61896 (CsbHLH121), ESR36836 (CcbHLH121), and XP_028243911 (GsbHLH121).

### Slide 5
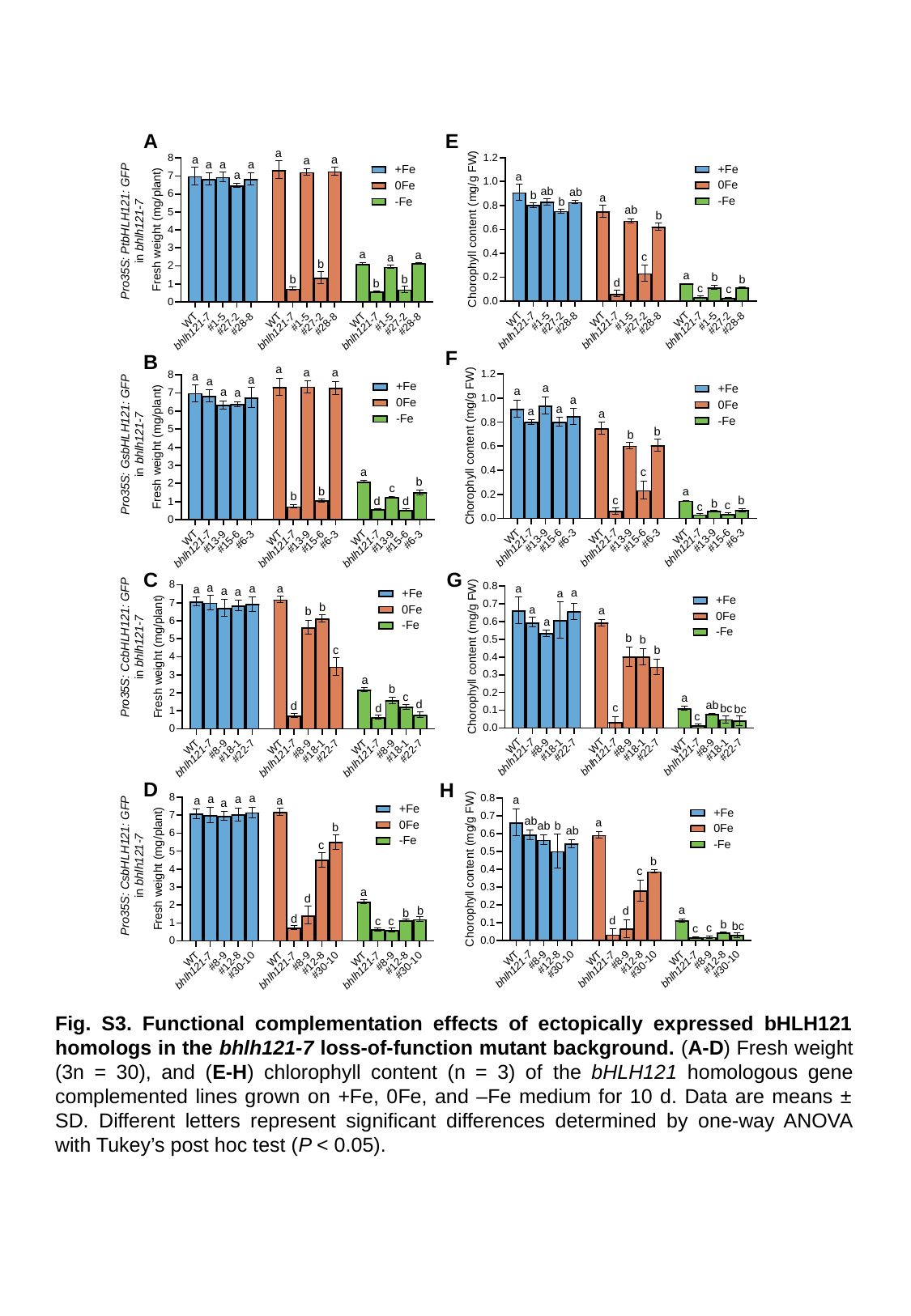

A
E
Pro35S: PtbHLH121: GFP
in bhlh121-7
F
B
Pro35S: GsbHLH121: GFP
in bhlh121-7
G
C
Pro35S: CcbHLH121: GFP
in bhlh121-7
D
H
Pro35S: CsbHLH121: GFP
in bhlh121-7
Fig. S3. Functional complementation effects of ectopically expressed bHLH121 homologs in the bhlh121-7 loss-of-function mutant background. (A-D) Fresh weight (3n = 30), and (E-H) chlorophyll content (n = 3) of the bHLH121 homologous gene complemented lines grown on +Fe, 0Fe, and –Fe medium for 10 d. Data are means ± SD. Different letters represent significant differences determined by one-way ANOVA with Tukey’s post hoc test (P < 0.05).

### Slide 6
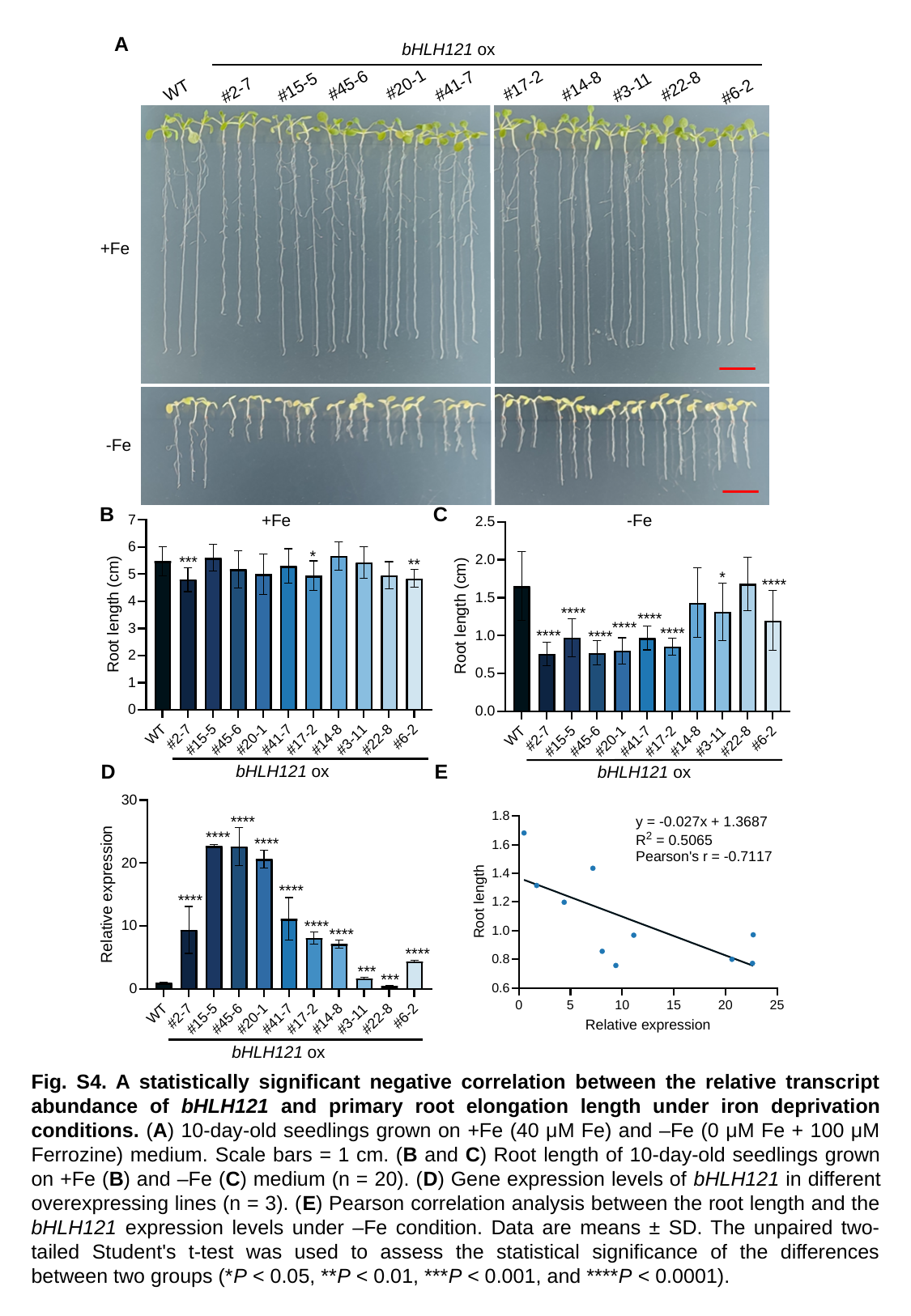

A
bHLH121 ox
#20-1
#45-6
#17-2
#41-7
#14-8
#22-8
#15-5
#3-11
WT
#2-7
#6-2
+Fe
-Fe
B
C
+Fe
-Fe
D
E
bHLH121 ox
bHLH121 ox
bHLH121 ox
Fig. S4. A statistically significant negative correlation between the relative transcript abundance of bHLH121 and primary root elongation length under iron deprivation conditions. (A) 10-day-old seedlings grown on +Fe (40 μM Fe) and –Fe (0 μM Fe + 100 μM Ferrozine) medium. Scale bars = 1 cm. (B and C) Root length of 10-day-old seedlings grown on +Fe (B) and –Fe (C) medium (n = 20). (D) Gene expression levels of bHLH121 in different overexpressing lines (n = 3). (E) Pearson correlation analysis between the root length and the bHLH121 expression levels under –Fe condition. Data are means ± SD. The unpaired two‐tailed Student's t‐test was used to assess the statistical significance of the differences between two groups (*P < 0.05, **P < 0.01, ***P < 0.001, and ****P < 0.0001).

### Slide 7
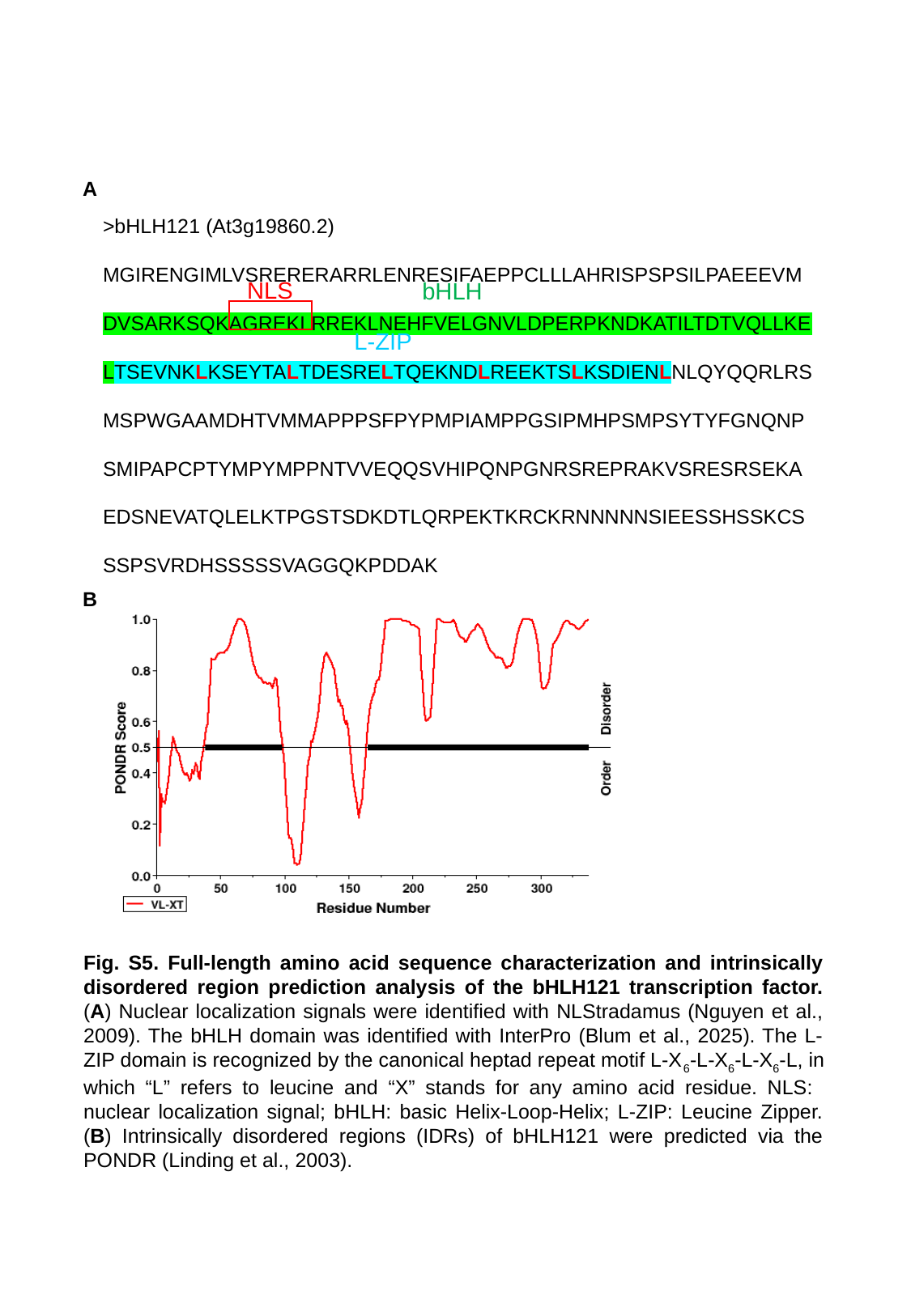

>bHLH121 (At3g19860.2)
MGIRENGIMLVSRERERARRLENRESIFAEPPCLLLAHRISPSPSILPAEEEVMDVSARKSQKAGREKLRREKLNEHFVELGNVLDPERPKNDKATILTDTVQLLKELTSEVNKLKSEYTALTDESRELTQEKNDLREEKTSLKSDIENLNLQYQQRLRS
MSPWGAAMDHTVMMAPPPSFPYPMPIAMPPGSIPMHPSMPSYTYFGNQNPSMIPAPCPTYMPYMPPNTVVEQQSVHIPQNPGNRSREPRAKVSRESRSEKAEDSNEVATQLELKTPGSTSDKDTLQRPEKTKRCKRNNNNNSIEESSHSSKCSSSPSVRDHSSSSSVAGGQKPDDAK
NLS
bHLH
L-ZIP
A
B
Fig. S5. Full-length amino acid sequence characterization and intrinsically disordered region prediction analysis of the bHLH121 transcription factor. (A) Nuclear localization signals were identified with NLStradamus (Nguyen et al., 2009). The bHLH domain was identified with InterPro (Blum et al., 2025). The L-ZIP domain is recognized by the canonical heptad repeat motif L-X_6_-L-X_6_-L-X_6_-L, in which “L” refers to leucine and “X” stands for any amino acid residue. NLS: nuclear localization signal; bHLH: basic Helix-Loop-Helix; L-ZIP: Leucine Zipper. (B) Intrinsically disordered regions (IDRs) of bHLH121 were predicted via the PONDR (Linding et al., 2003).

### Slide 8
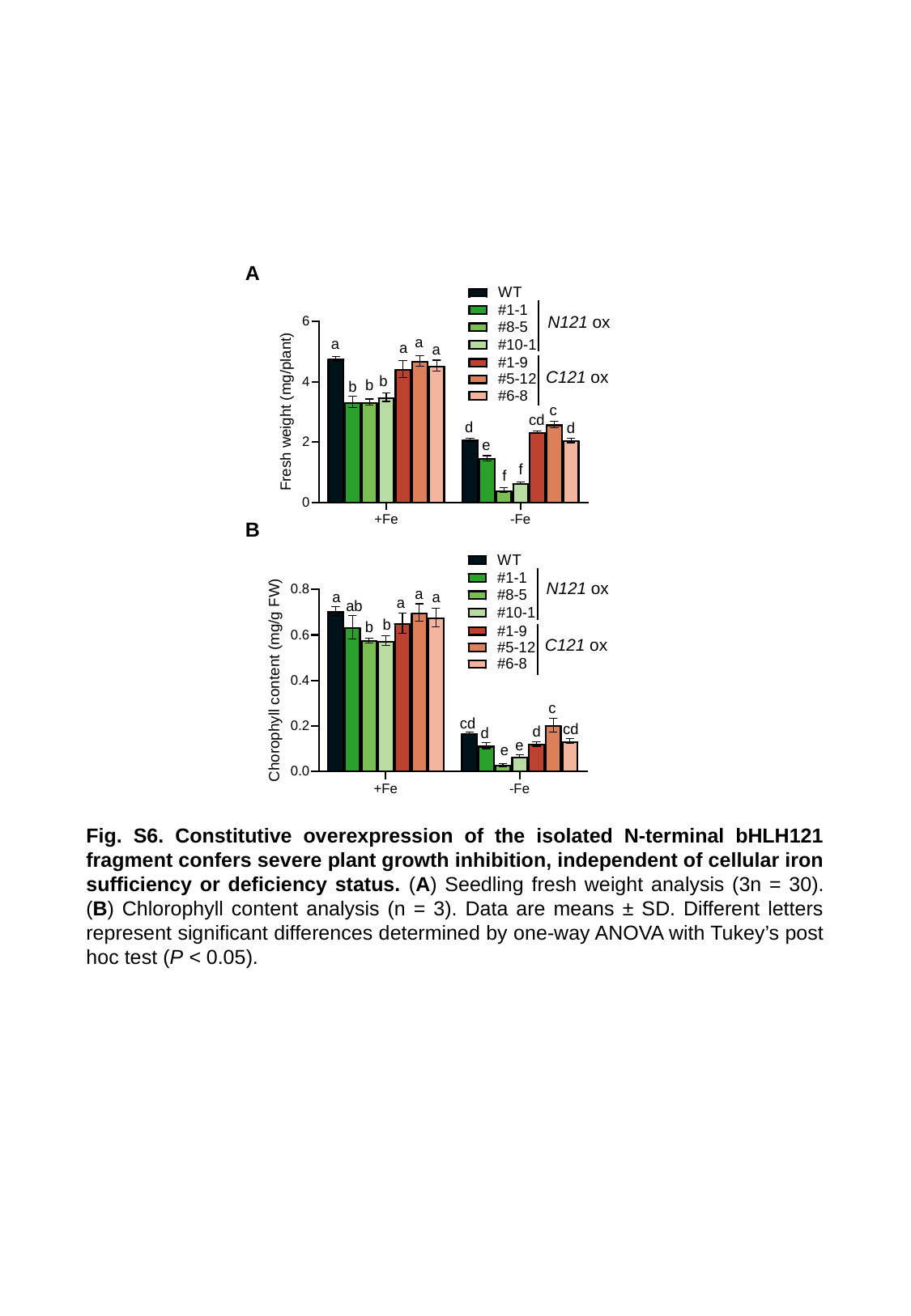

A
N121 ox
C121 ox
B
N121 ox
C121 ox
Fig. S6. Constitutive overexpression of the isolated N-terminal bHLH121 fragment confers severe plant growth inhibition, independent of cellular iron sufficiency or deficiency status. (A) Seedling fresh weight analysis (3n = 30). (B) Chlorophyll content analysis (n = 3). Data are means ± SD. Different letters represent significant differences determined by one-way ANOVA with Tukey’s post hoc test (P < 0.05).

### Slide 9
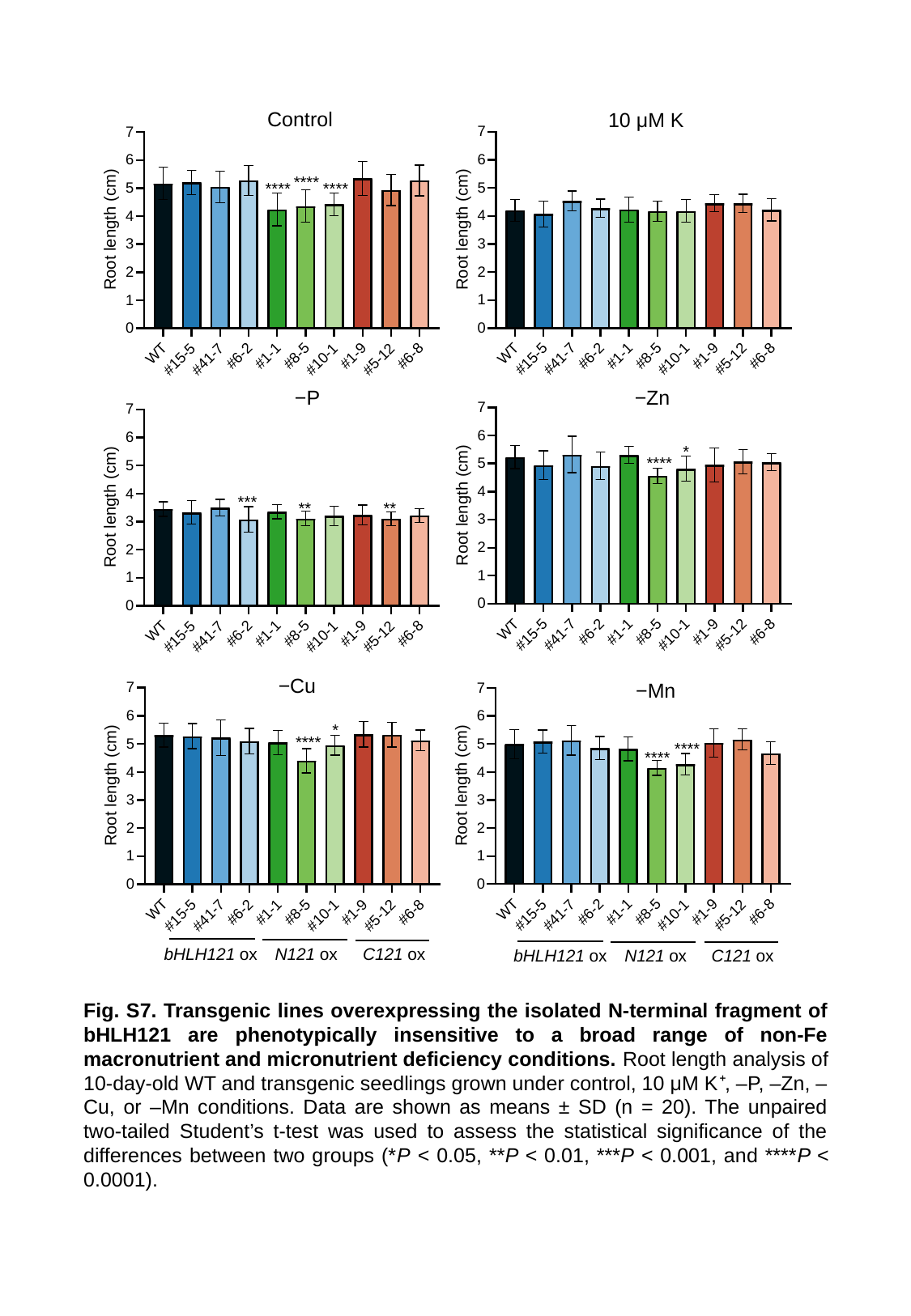

Control
10 μM K
−Zn
−P
−Cu
−Mn
bHLH121 ox
N121 ox
C121 ox
bHLH121 ox
N121 ox
C121 ox
Fig. S7. Transgenic lines overexpressing the isolated N-terminal fragment of bHLH121 are phenotypically insensitive to a broad range of non-Fe macronutrient and micronutrient deficiency conditions. Root length analysis of 10-day-old WT and transgenic seedlings grown under control, 10 μM K⁺, –P, –Zn, –Cu, or –Mn conditions. Data are shown as means ± SD (n = 20). The unpaired two‐tailed Student’s t‐test was used to assess the statistical significance of the differences between two groups (*P < 0.05, **P < 0.01, ***P < 0.001, and ****P < 0.0001).

### Slide 10
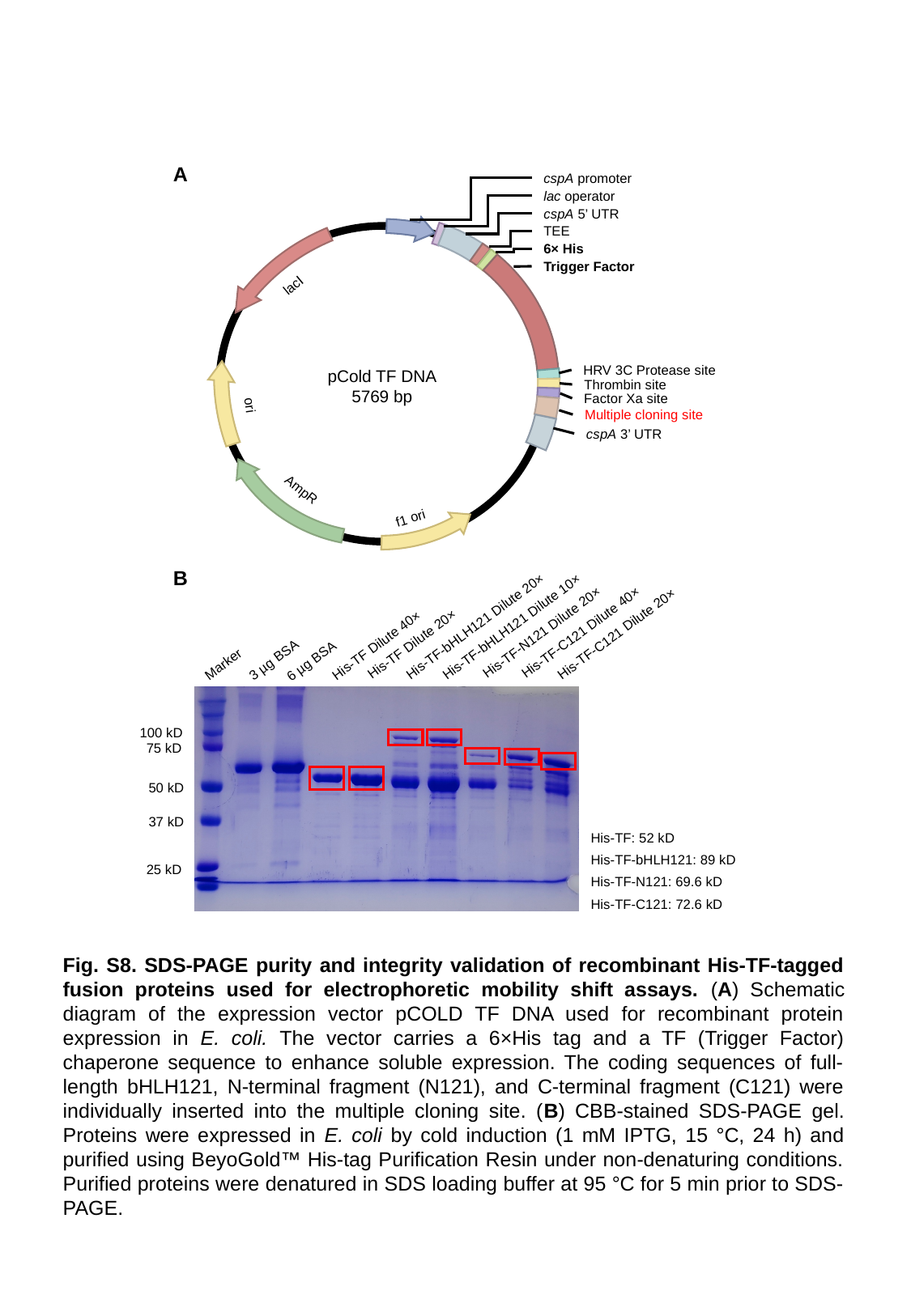

A
cspA promoter
lac operator
cspA 5’ UTR
TEE
6× His
Trigger Factor
lacⅠ
HRV 3C Protease site
pCold TF DNA
5769 bp
Thrombin site
Factor Xa site
ori
Multiple cloning site
cspA 3’ UTR
AmpR
f1 ori
B
His-TF-N121 Dilute 20×
His-TF Dilute 20×
His-TF-C121 Dilute 40×
His-TF-bHLH121 Dilute 20×
His-TF-bHLH121 Dilute 10×
His-TF-C121 Dilute 20×
His-TF Dilute 40×
3 µg BSA
6 µg BSA
Marker
100 kD
75 kD
50 kD
37 kD
His-TF: 52 kD
His-TF-bHLH121: 89 kD
25 kD
His-TF-N121: 69.6 kD
His-TF-C121: 72.6 kD
Fig. S8. SDS-PAGE purity and integrity validation of recombinant His-TF-tagged fusion proteins used for electrophoretic mobility shift assays. (A) Schematic diagram of the expression vector pCOLD TF DNA used for recombinant protein expression in E. coli. The vector carries a 6×His tag and a TF (Trigger Factor) chaperone sequence to enhance soluble expression. The coding sequences of full-length bHLH121, N-terminal fragment (N121), and C-terminal fragment (C121) were individually inserted into the multiple cloning site. (B) CBB-stained SDS-PAGE gel. Proteins were expressed in E. coli by cold induction (1 mM IPTG, 15 °C, 24 h) and purified using BeyoGold™ His-tag Purification Resin under non-denaturing conditions. Purified proteins were denatured in SDS loading buffer at 95 °C for 5 min prior to SDS-PAGE.
